# Covalent Drug Binding in Live Cells Monitored by Mid-IR Quantum Cascade Laser Spectroscopy: Photoactive Yellow Protein as a Model System

**DOI:** 10.1101/2025.08.15.670201

**Authors:** Srijit Mukherjee, Steven D. E. Fried, Nathalie Y. Hong, Nahal Bagheri, Jacek Kozuch, Irimpan I. Mathews, Jacob M. Kirsh, Steven G. Boxer

## Abstract

The detection of drug-target interactions in live cells enables analysis of therapeutic compounds in a native cellular environment. Recent advances in spectroscopy and molecular biology have facilitated the development of genetically encoded vibrational probes like nitriles that can sensitively report on molecular interactions. Nitriles are powerful tools for measuring electrostatic environments within condensed media like proteins, but such measurements in live cells have been hindered by low signal-to-noise ratios. In this study, we design a spectrometer based on a double-beam quantum cascade laser (QCL)-based transmission infrared (IR) source with balanced detection that can significantly enhance sensitivity to nitrile vibrational probes embedded in proteins within cells compared to a conventional FTIR spectrometer. Using this approach, we detect small-molecule binding in *E. coli*, with particular focus on the interaction between para-coumaric acid (pCA) and nitrile-incorporated photoactive yellow protein (PYP). This system effectively serves as a model for investigating covalent drug binding in a cellular environment. Notably, we observe large spectral shifts of up to 15 cm^−1^ for nitriles embedded in PYP between the unbound and drug-bound states directly within bacteria, in agreement with observations for purified proteins. Such large spectral shifts are ascribed to the changes in the hydrogen-bonding environment around the local environment of nitriles, accurately modeled through high-level molecular dynamics simulations using the AMOEBA force field. Our findings underscore the QCL spectrometer’s ability to enhance sensitivity for monitoring drug-protein interactions, offering new opportunities for advanced methodologies in drug development and biochemical research.

Authors are required to submit a graphic entry for the Table of Contents (TOC) that, in conjunction with the manuscript title, should give the reader a representative idea of one of the following: A key structure, reaction, equation, concept, or theorem, etc., that is discussed in the manuscript. Consult the journal’s Instructions for Authors for TOC graphic specifications.

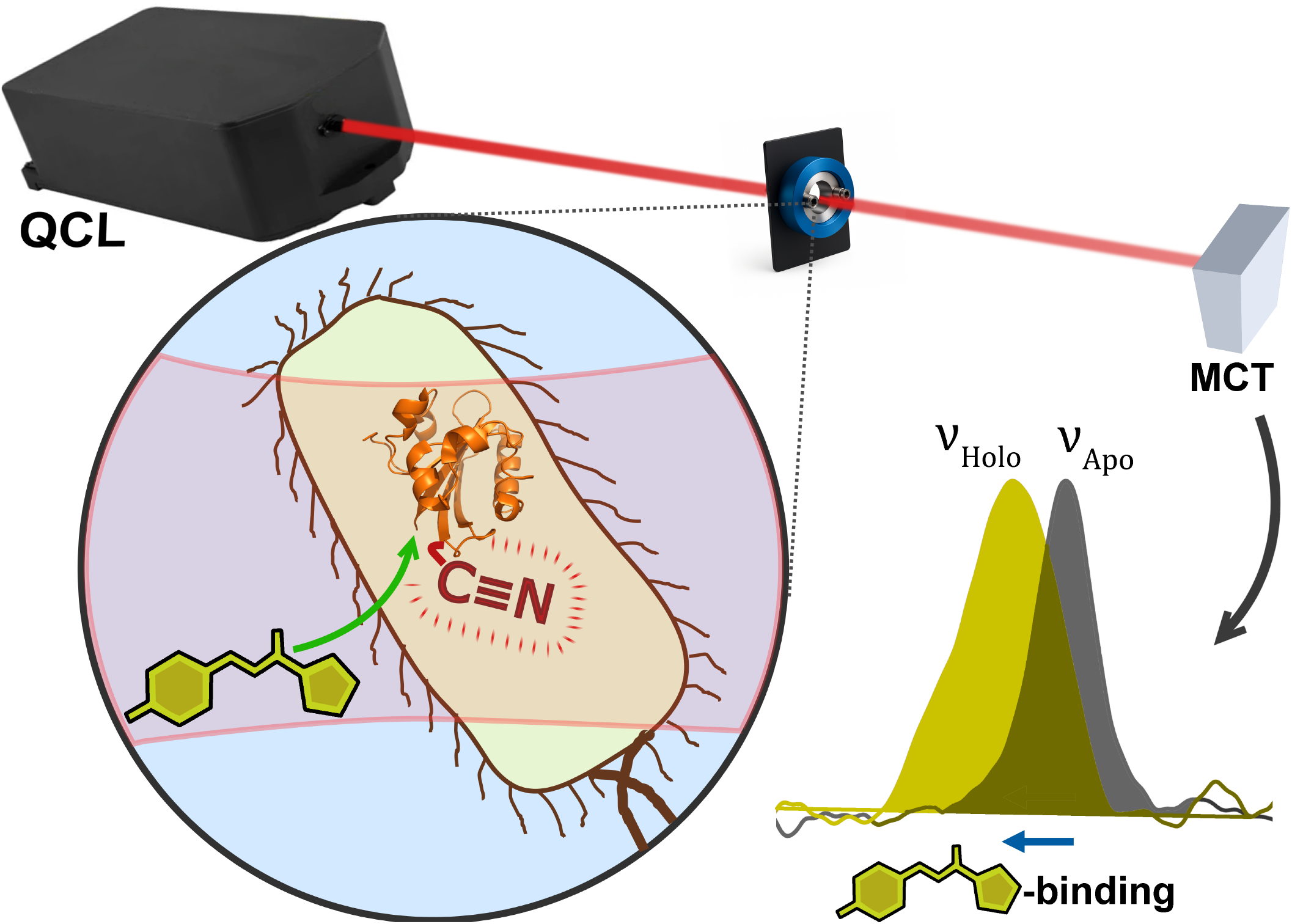

## Introduction

Detection and quantification of drug binding is one of the most sought-after applications of analytical chemistry to human health and comprises a multi-billion-dollar industry. While many drug binding techniques rely upon separative methods or highly perturbative techniques (e.g., calorimetry, surface plasmon resonance, mass spectrometry) that can only be carried out *in vitro*^1,2,3^ there is a decisive advantage to being able to directly probe drug-protein interactions in living cells. Measurements of drug binding in live cells can capture the native environment of the target protein of interest and account for the effects of competing variables that might limit drug binding to a target protein in a cell, including metabolic degradation, trafficking to the correct compartment, or competition of off-target sites for binding to the drug of interest.^4^

While cell-based assays remain essential for assessing efficacy and toxicity—the ultimate goals of drug discovery— they often rely on downstream, *indirect* readouts. ^5^ In contrast, direct measurements of drug binding to the protein of interest can provide mechanistic insights into target engagement and binding mode, offering valuable complementary information. ^6,7^ The majority of such live-cell measurements make use of fluorescence-based assays based on fluorescence polarization,^8^ Förster Resonance Energy Transfer (FRET),^9^ fluorescence lifetime imaging (FLIM),^10^ fluorescence thermal shift assays (FTSA)^11^ and single molecule fluorescence.^12^ Yet, such assays are disadvantaged by limited applicability for non-fluorescent drug systems and generally require labeling by bulky fluorescent groups such as dyes or fluorescent proteins that have the potential to be highly perturbative to the system of interest.^13,14^

Vibrational probes by contrast can serve as minimally perturbative reporters of their immediate environment, demonstrating frequency shifts that are physically interpretable in terms of environmental electrostatics^15,16,17^ molecular conformation,^18^ or hydrogen bonding.^19,20,21^ Many common vibrational probes like nitriles (C≡N) are intrinsic to drugs themselves^22^ or can be engineered selectively onto the target protein scaffold via amber suppression^23,24^ and can thus serve as effective reporters of drug-protein interactions. Nitriles are of particular interest for this purpose due to their relatively strong absorption in a clean region of the infrared spectrum. They have been used as molecular probes in various contexts, including proteins,^21-25^ lipid membranes,^26^ surfaces,^27^ microdroplets^28^ and metal-organic frameworks.^17^

*In cellulo* spectroscopic analysis of site-specifically labeled vibrational probes represents a relatively unexplored frontier—one that remains limited by both sensitivity to low concentrations and interference from cellular background. Most efforts in live-cell transmission IR spectroscopy focus on lineshape differences in the amide I and II regions, rather than the detection of specific vibrational frequency shifts of labeled sites.^29,30,^31,32 Despite these difficulties, a few recent and notable studies have demonstrated progress, typically using attenuated total reflectance (ATR)-based techniques to focus on strong absorbing oscillators such as μ-CO in hydrogenases in whole *E. coli*^33^ or flavin carbonyls as they photo-react in human cell lines.^34^ Another recent study stemming from Heberle and co-workers has made use of high-power quantum cascade lasers (QCLs) for time-resolved detection of cysteine protonation dynamics in halorhodopsin in live *E. coli*.^35^ QCLs pose a significant advantage as a mid-IR light source in that their power at a given frequency is orders of magnitude higher than conventional globar sources, enabling substantially longer sample pathlengths.^36^ For example, QCLs emit tunable, narrow-bandwidth mid-infrared light that has advanced discrete frequency mid-infrared imaging for label-free, high-resolution histopathological analysis.^37^ Lendl and coworkers have demonstrated the remarkable capabilities of external cavity, mid-IR QCLs for biomolecular spectroscopy,^38,39,40^ especially how the use of balanced detection with a double-beam approach can compensate for much of the noise inherent in QCL radiation and enhance the sensitivity of transmission IR spectroscopy measurements for the amide region of proteins in aqueous buffer.^41,42^

Here we extend the capabilities of QCL-based vibrational spectroscopy with balanced detection for monitoring genetically encoded nitrile vibrational probes in live cells as molecular reporters of protein-small molecule binding. We first demonstrate a significant sensitivity improvement, up to five-fold, over conventional FTIR spectroscopy for detection of aromatic nitriles due to longer pathlengths for transmission measurements accessible to the QCL setup. We then use the QCL spectrometer to detect small molecule binding to proteins—a key objective in drug discovery—using nitrile frequency shifts as direct indicators of fine protein structural perturbations caused by binding events in live cells.

To model covalent drug-binding events, we present an example based on photoactive yellow protein (PYP) from *Halorhodospira halophila. Apo*-PYP, as expressed in cells, forms a covalent complex with the chromophore, p-coumaric acid (pCA), a process that is analogous to covalent drug binding. PYP is a highly soluble globular protein from the PAS domain superfamily.^43^ While not a drug target in the conventional sense, it shares chemical characteristics with common covalent drug targets, especially those with reactive cysteines.^44,45,46,47^ Moreover, PYP’s potential as an optogenetic system has led to considerable interest in measuring its interactions with its chromophore and other binding partners in living cells.^43,48^ While a FRET-based assay using a fusion of a blue fluorescent protein and *apo*-PYP has previously enabled an *in cellulo* binding study,^49^ we note that a vibrational spectroscopy approach without the use of a fusion protein construct is much more generalizable including non-photoactive compounds involved in other systems. Our spectroscopic analyses of the nitrile frequency shifts using this assay, supported by detailed structural studies including X-ray crystallography and high-level AMOEBA molecular dynamics (MD) simulations, indicate subtle changes in hydrogen bonding interactions involving the embedded nitriles within the protein upon drug incorporation. These findings underscore the utility of this assay for measuring drug binding in live cells at the molecular level using QCL-based IR spectroscopy frequency shifts.

## Results

### Design and use of the dual-beam QCL-spectrometer for transmission IR measurements

We designed a custom double-beam infrared spectrometer, employing an external-cavity quantum cascade laser (EC-QCL) as the mid-IR radiation source (MIRcat-QT-2000 with M2048-P, Daylight Solutions, San Diego, CA, USA), enabling spectral acquisition within the 2000-2300 cm^−1^ range (Figs. S1, S2) using an optical layout inspired from the work of Akghar et al.^41^ The instrument utilizes a balanced detection scheme, where the incident laser beam is split into reference and sample paths, and subsequently focused onto a thermoelectrically cooled HgCdTe (MCT) detector (VIGO Photonics, Poland). The optical diagram is presented in Fig. 1 with calibration information in Section S1, Figs. S3, S4. A high-speed SR865A lock-in amplifier (Stanford Research Systems, Sunnyvale, CA, USA) and custom MATLAB routines were used for data acquisition and signal processing, including phase correction, Savitsky-Golay filtering, and Fourier filtering.^38^ Absorbance measurements were derived from the ratio of beam intensities taken from reference and balanced channel voltage outputs, effectively mitigating fluctuations, primarily driven by the QCL source, using the balanced detection scheme. Two identical liquid sample cells equipped with CaF_2_ windows and Teflon spacers (50-250 µm; Pike Technologies, Fitchburg, WI, USA) were mounted on machined sample holders, serving as the sample and blank sample cells.

**Figure 1.**
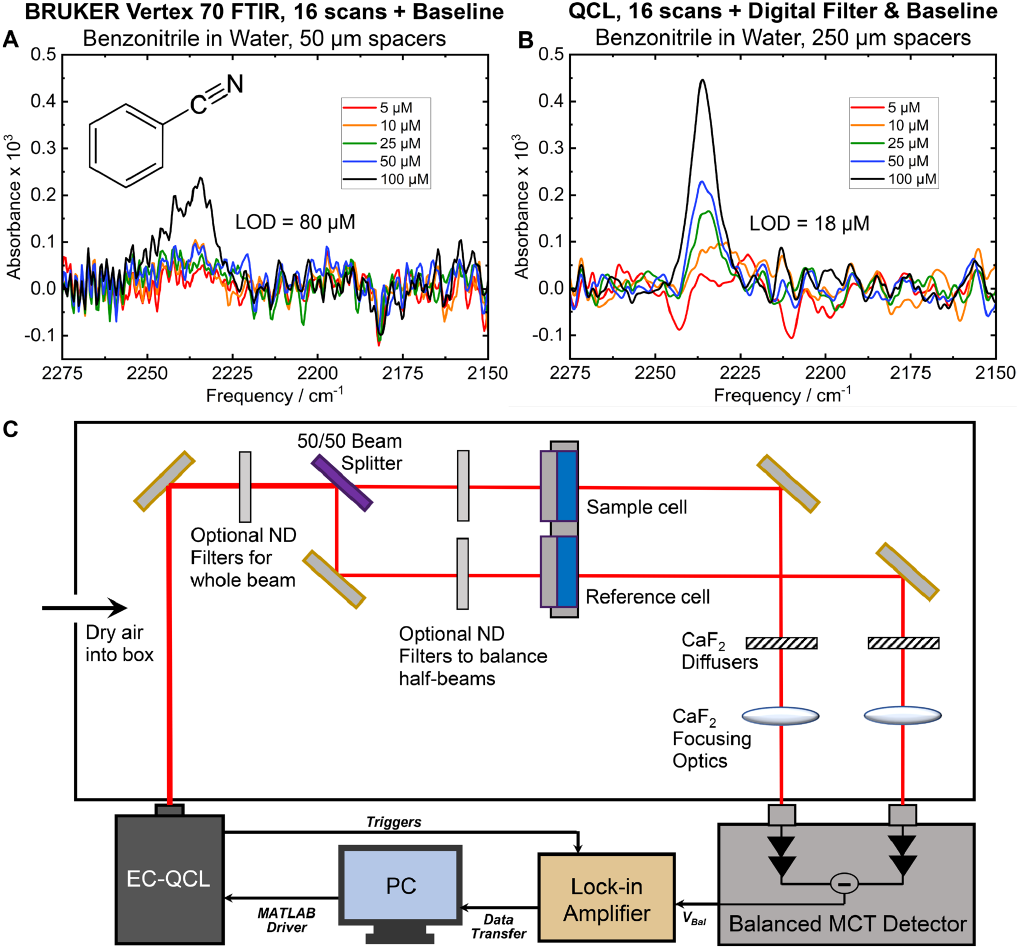
Comparison of detection limits for benzonitrile using (A) a conventional Bruker Vertex-70 FTIR spectrometer and (B) a QCL-based spectrometer. The FTIR, limited by aqueous sample transmission with 50 µm spacers, achieves a limit of detection (LOD) of 80 µM. The QCL-based approach improves the LOD to 18 µM, offering a ∼5-fold enhancement in sensitivity for IR measurements in the nitrile stretch region compared to high-end FTIR systems. (C) Block diagram of double-beam QCL spectrometer with lock-in detection for balanced absorption measurement.

Utilizing benzonitrile as a model nitrile probe in water in the µM range, we compared the limit of detection for the spectrometer against a conventional Bruker Vertex 70 FTIR spectrometer (Figs. S5-S9, Tables S1-S3). We also document in the SI the increased sensitivity of the QCL for other useful vibrational probes including alkyne (C≡C), azides (N=N^+^=N), and carbon-deuterium (C–D) stretches (Figs. S10-S12, Table S4). A notable challenge for IR spectra of aqueous samples in this spectral window is restricted transmission due to water’s combination mode centered at approximately 2100 cm^−1^.^50^ Consequently, this imposes a limit for the maximum spacer length that can be used in the Bruker Vertex 70 at ∼50 µm leading to a limit of detection (LOD) for benzonitrile at 80 µM for the nitrile stretch centered at ∼2235 cm^-1^. In contrast, the balanced QCL-based detection using 250 µm spacers significantly enhances the experimental LOD, achieving a value of 18 µM. This QCL-based method demonstrates an approximately five-fold increase in sensitivity for IR measurements within the nitrile stretch region, outperforming the conventional FTIR spectrometer and enabling detection of low concentrations of nitriles in aqueous environments such as proteins or molecules labeled with nitriles directly within cells (Fig. 1, Table S5).

### Incorporation and detection of nitrile-labeled proteins in live bacterial cells

We evaluated the spectrometer’s ability to detect nitrile stretches within *E. coli* cells using amber suppression machinery to introduce ortho-cyanophenylalanine (oCNF) residues into target proteins of interest. Our primary focus was on *apo*-PYP as a model system for the equivalent of covalent drug binding. We used site-specific labeling to produce four *apo*-PYP variants: F28oCNF, F62oCNF, F92oCNF, and F96oCNF. As an additional test of visualizing nitriles in live cells we also incorporated a nitrile into superfolder green fluorescent protein (sfGFP), where the chromophore-forming tyrosine Y66 was replaced with oCNF. See Supplementary Information Section S2 for further details on sample preparation.

The variable proximities of the four nitrile probes on PYP to the pCA binding site allowed us to observe locally specific information on subtle structure perturbations when activated pCA binds both in solution and in living cells. Each nitrile-containing PYP variant was previously characterized in the *holo*-form (covalently attached to pCA) via X-ray crystallography to high-resolution (<1.2 Å), where the nitriles adopt a single, well-defined orientation at each site (Fig. S13). ^23^ F28oCNF (PDB: 7SPX) interacts with one hydrogen bond donor, a water molecule (3.2 Å heavy atom distance) and is located ∼15 Å (ring-to-ring distance) from pCA, while F92oCNF (PDB: 7SPV) engages with two donors (2.9 and 3.2 Å for T90 and a water, respectively) and is positioned ∼12 Å from the pCA chromophore. In contrast, F62oCNF (PDB: 7SPW) and F96oCNF (PDB: 7SJJ) are situated in hydrophobic environments, with only carbon atoms within 3.5 Å of the nitrile nitrogen, and they are located ∼10 Å and ∼5 Å from the pCA chromophore, respectively. As part of this study, we also determined for the first time a high-resolution X-ray crystal structure of the *apo* form of PYP (without the nitrile labels; PDB: 9O8V) and observed minimal deviation from the *holo* form, both with and without a nitrile incorporated (Fig. 2, SI Section S3 and Table S6). The absence of the chromophore and nitrile labels did not result in any major conformational changes in the protein structure, except for a lack of density in the disordered loop region (residues 16–20), which is located more than 20 Å from the chromophore pocket (Fig. S14).

**Figure 2.**
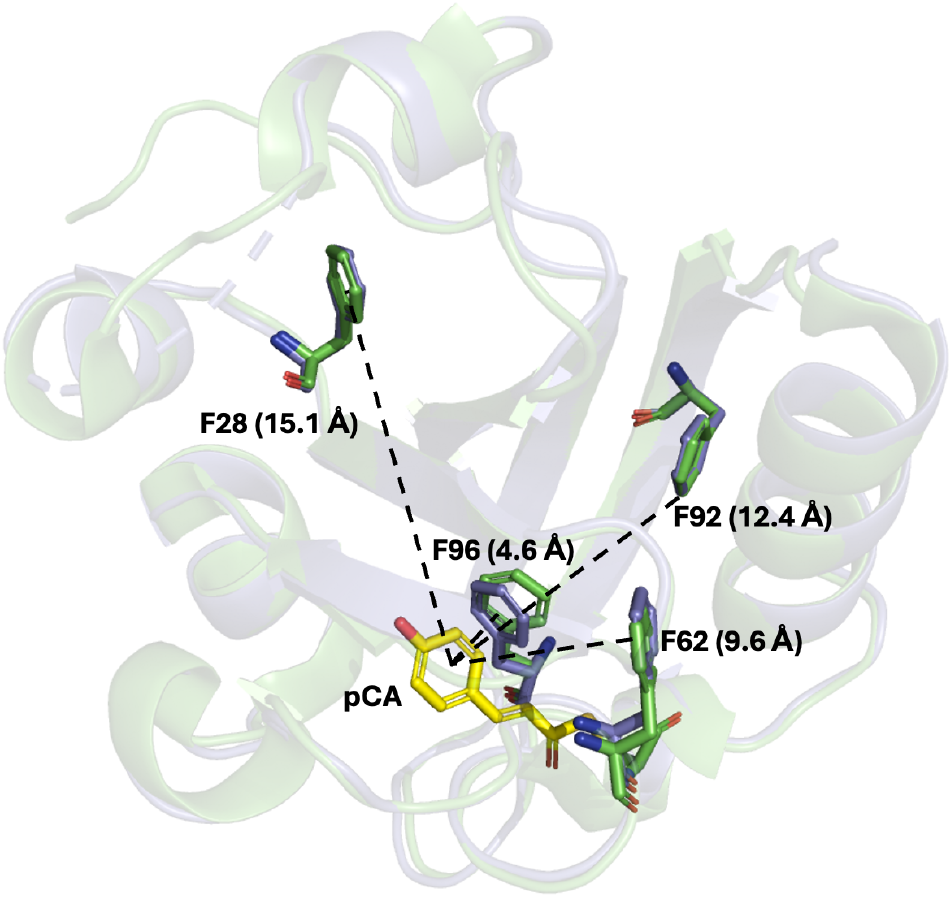
Strategy to study covalent drug binding using photoactive yellow protein (PYP) and nitrile vibrational spectroscopy. oCNF incorporation via amber suppression allows site-specific placement of nitrile reporters at F28, F62, F92, and F96. *Holo* (Green, PDB: 1NWZ) and *apo* (Grey, PDB: 9O8V) PYP crystal structures superimposed with the modified Phe residues highlighted.

Utilizing these variants and the sensitive QCL spectrometer, we measured the nitrile absorption spectra in a concentrated suspension of bacterial cells (OD_600_ = 0.3 when diluted 1000x) expressing individually nitrile-modified Phe variants of *apo*-PYP. Cells were washed three times with minimal media to remove excess unincorporated oCNF (See SI Section S2). A culture expressing unmodified *apo*-PYP (no nitrile) was used as a blank for balanced detection. We could clearly note the nitrile frequencies: for F28oCNF at 2230.8 cm^-1^, for F62oCNF at 2227.5 cm^-1^, for F92oCNF at 2226.4 cm^-1^, and for F96oCNF at 2227.6 cm^-1^ (Fig. 3). While similar measurements were attempted using the Bruker 70 FTIR spectrometer with 50 µm spacers (SI Section S4), the low signal-to-noise ratio hindered reliable detection of these peaks (Fig. S15, Table S7). For Y66oCNF sfGFP, the nitrile stretch was also clearly visible at 2225.8 cm^−1^ in cells using the QCL spectrometer (Fig. S16).

**Figure 3.**
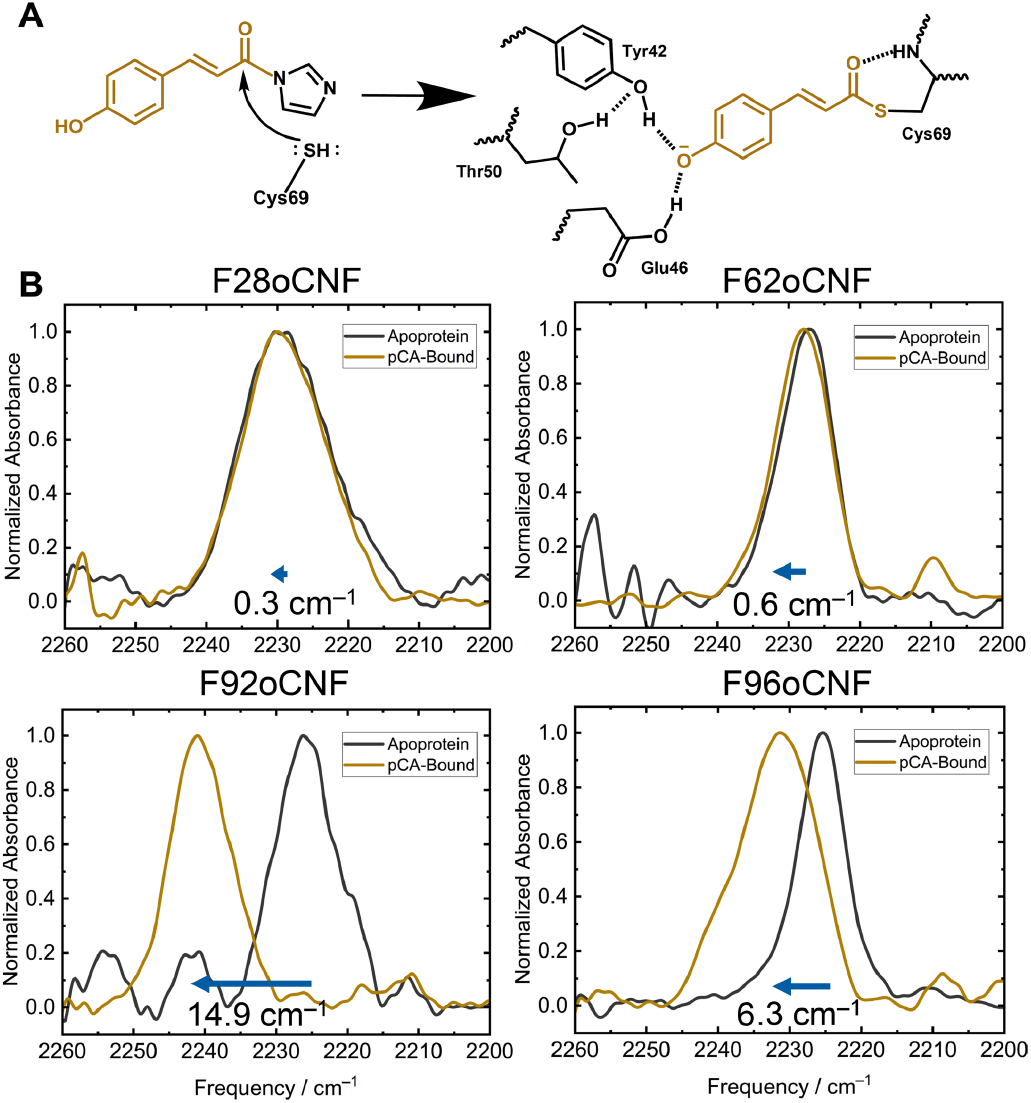
(A) Incorporation of the chromophore p-coumaric acid (pCA) into PYP proceeds by nucleophilic attack at C69. The form of pCA added to the cells is an activated form with a reactive, electrophilic “warhead” that derives from a water-free reaction of pCA with carbonyldiimidazole. (B) Nitrile vibrational spectra collected in bacteria using the QCL for the F28oCNF, F62oCNF, F92oCNF, and F96oCNF variants show variable nitrile vibrational frequency shifts in response to pCA incorporation, with shifts of +0.3, +0.6, +14.9, and +6.3 cm^-1^, respectively (analyzed by peak fitting).

In all cases, the *in cellulo* nitrile frequencies were distinct from that of free oCNF in aqueous media (2232.3 cm^−1^ in water; Fig. S16). Furthermore, when cells expressing nitrile-containing proteins were lysed and purified by affinity column chromatography, the nitrile stretching frequencies of the isolated proteins matched those observed in cells (Figs. S16-S18). However, when *apo*-PYP nitrile frequencies were compared to previous measurements conducted on the purified *holo*-PYP (with pCA covalently attached), significant differences in peak frequency were noted for variants F92oCNF and F96oCNF (Tables S8-S9). In contrast, the frequencies for F28oCNF and F62oCNF were similar regardless of chromophore incorporation. These findings indicated that some of the nitrile environments in the protein depend on pCA binding, prompting further investigations described below.

### Observation of nitrile frequency shifts on pCA-binding from the QCL-based IR measurement

We next incubated the PYP-expressing bacteria with a ∼100-fold molar excess of pCA chemically activated with an acyl imidazolide electrophilic warhead (SI Section S5, Figs. S19-S25).^23,24,51,52^ The nitrile stretch measured by the QCL in bacteria expressing F28oCNF or F62oCNF PYP exhibited negligible shifts after activated pCA incubation, while those expressing F92oCNF and F96oCNF displayed large blue shifts of +14.9 cm^−1^ and + 6.3 cm^−1^, respectively, when compared to the *apo* samples. These nitrile frequencies align closely with those of the previously reported, isolated *holo* F92oCNF and F96oCNF proteins.^23^ Subsequent purification of these activated pCA-incubated cells yielded a characteristic yellow protein with a UV-Vis absorbance maximum at 445 nm, which along with mass spectrometry (SI Section S6, Fig. S26) confirmed chromophore incorporation through the cell wall and membrane and into the *apo*-protein. Additionally, the nitrile frequencies for these purified proteins closely matched the frequencies of both the previously reported purified *holo*-PYP^23^ and the QCL-measured *in cellulo* nitrile frequencies of *holo*-PYP (Tables S8-S11).

As a control, we introduced a C69A substitution in the F96oCNF variant (F96oCNF was selected due its ∼6 cm^−1^ blue shift between the *holo* and *apo* forms). The C69A mutation thereby disrupted the covalent linkage site for pCA in the F96oCNF-PYP variant. Even after activated pCA incubation, the nitrile frequency in this variant remained similar to that of the *apo*-F96oCNF. This confirms that the C69 thiol group is indeed responsible for pCA binding including in the cellular environment, and the covalent linkage of pCA in the binding site accounts for the observed frequency shifts (Fig. S18, Table S12).

### Consequences of drug binding to *apo*-PYP—structural analysis and MD simulations

We next sought to better understand the origin of the large frequency shifts on pCA incorporation for F92oCNF and F96oCNF PYP. As demonstrated by our group^23^ and others^19^, nitrile vibrational frequency shifts are affected by intermolecular interactions— electrostatic stabilization typically causes red shifts, while hydrogen bonding can induce blue shifts with respect to an unperturbed nitrile stretch.^20^ Particularly when the concentration of the vibrational probe cannot be obtained with high accuracy,^23**\***^ interpreting a spectroscopically observed frequency shift requires structural knowledge of the local electric fields and hydrogen-bonding populations. We investigated these influences on nitrile frequency shifts in PYP using molecular dynamics simulations.

We first ran 100-ns MD simulations of the *apo*-form using the AMBER99SB-ILDN forcefield.^53^ This revealed no global conformational changes, whether starting from the *apo* crystallographic pose first reported here (Fig. 2) or the *holo*-crystallographic pose 1NWZ but with pCA removed (Fig. S27). As we have shown in separate work, fixed-charge MD simulations based on force-fields such as AMBER often cannot accurately recapitulate the electrostatic environment around nitrile probes in proteins, thus for these analyses higher-level simulations were carried out using the AMOEBA09 forcefield (Fig 4).^54,55^ Four replicate AMOEBA simulations of 25 ns each were performed using the F92oCNF (7SPV) and F96oCNF (7SJJ) PYP structures with chromophores removed as starting points,^23^ again showing no global conformational changes (Fig. S28, details in SI Section S7).^56^ These simulations revealed that for *apo*F92oCNF, the average electric field projected on the nitrile bond was significantly reduced to –44 MV/cm, compared to the –67 MV/cm reported by Kirsh et al. for the *holo*-form.^24^ *Apo*-F96oCNF showed only a slightly smaller average field, –23 MV/cm compared to –26 MV/cm reported for the *holo*form. ^24^ A more stabilizing field on the nitriles in the *holo*form would typically coincide with a red-shift upon pCA binding. However, the presence of hydrogen bonds as is possible for both nitriles^24^ obfuscates a purely electrostatic interpretation of the observed frequency differences, in contrast to the transition dipole moment approach used by Weaver et al.^23^ and further discussed in ref 24.

**Figure 4.**
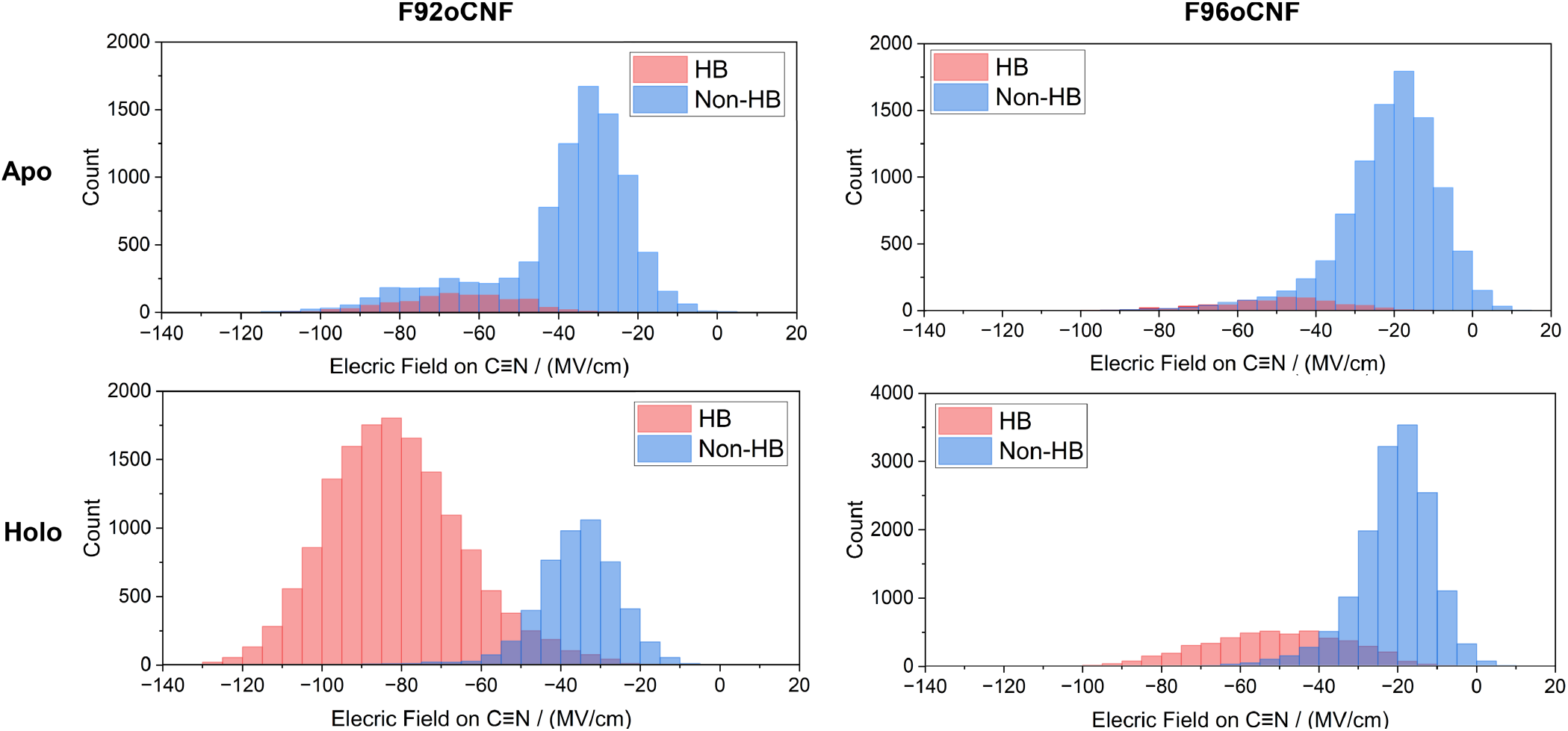
Local electrostatic environment around the nitrile group revealed by AMOEBA molecular dynamics simulations. Simulations were run for four replicates of 25 ns each. The distribution of electric fields experienced by the nitrile is shown, separated by whether the nitrile was engaged in hydrogen bonding. Hydrogen bonding was defined using a heavy-atom nitrile–H-bond donor–acceptor distance cutoff of 4.0 Å and a donor–acceptor–hydrogen angle cutoff of 30°. F92oCNF exhibits two distinct electrostatic states: a non-hydrogen-bonded (low-field) state and a hydrogen-bonded (high-field) state, which significantly shift in population upon pCA binding—correlating with a large ∼15 cm^−1^ blue shift in the nitrile stretch. F96oCNF shows a similar but smaller increase in the hydrogen-bonded population, consistent with the more modest 6.3 cm^−1^ blue shift observed upon pCA binding. Notably, the simulated electric field distributions are significantly broader than the experimentally observed linewidths, suggesting additional structural constraints or dynamic averaging in the actual protein environment.

For F92oCNF (Fig. 4), we clearly observed two distinct electric field states projected on the nitrile bond: a high-field state typically corresponding to a hydrogen-bonded state (primarily with T90 side chain), and a significantly lower-field state corresponding to a non-hydrogen-bonded configuration. Distinct hydrogen-bonding populations agree with previous low-temperature FTIR experiments of the *holo* form that slow the rate of exchange between the two populations (SI Section S8, Fig. S29).^24^ They also align with previous MD observations of a hydrogen-bonding and a non-hydrogen-bonding state in F92oCNF when the chromophore was present, where much of the simulation trajectory indicated a hydrogen-bonded state. ^24^ The current simulations in the absence of the chromophore significantly favor decreased hydrogen bonding of the nitrile to T90 in *apo*-F92oCNF, consistent with the IR measurements showing a blue shift upon chromophore incorporation. Additionally, AMOEBA simulations reveal a small increase in hydrogen-bonded population (nitrile-to-solvent) for the *apo*-F96oCNF variant upon pCA binding. We note it may be more difficult to reliably interpret simulation results in the vicinity of the chromophore pocket given the more dynamic solvation environment, particularly when pCA is absent and the pocket is solvent-exposed (Fig. 2). However, the small but definite change in solvent hydrogen bonding to the nitrile in F96oCNF presents a reasonable explanation for the smaller blue shift undergone by the nitrile upon pCA binding.

## Discussion

The IR frequency of a unique vibrational mode such as a nitrile is highly sensitive to changes in its local environment. This makes frequency shifts upon drug binding an indispensable tool for studying drug–protein interactions. However, the low sensitivity of this method *in cellulo* means it has rarely been used to date, instead opting for more traditional fluorescence-based assays. Here, we address this limitation by developing a QCL-based infrared spectrometer capable of sensitively detecting site-specific vibrational shifts in proteins bearing embedded nitrile probes, highlighting its utility for monitoring protein-drug interactions even within live-cell environments. Compared to advanced Raman techniques such as stimulated Raman spectroscopy (SRS), which can also be applied *in cellulo*,^57^ linear QCL spectroscopy leveraging long path lengths can still provide great sensitivity for bulk measurements.^58^ While methods like SRS offer high-resolution imaging and enhanced sensitivity when using vibrational probes conjugated to electronic chromophores (e.g., epr-SRS, Bon-FIRE), ^59^ the chemical interpretability of spectral intensities is not straightforward, and sensitivity is typically lower when using ordinary vibrational probes due to their inherently lower molecular cross-sections. (e.g., 10^5^ molecules/0.1 fL ≈ 1 mM for C=C bonds) ^60,61^

We present the first implementation of quantum cascade laser (QCL)-based balanced detection specifically optimized for the nitrile stretching region (2000–2300 cm^−1^), a spectrally transparent window with minimal interference from endogenous biomolecular absorption, enabling high-sensitivity detection in complex biological environments. Based on calculated protein expression levels, we estimate our working nitrile concentration within each cell to be approximately 140 μM (SI Section S2)—well above our determined 18 μM LOD for aromatic nitriles using the QCL and at a considerably higher SNR when compared to a traditional FTIR instrument (Fig. 3 vs. Fig. S15). This highlights the improved or at least comparable sensitivity of our approach relative to previous methodologies.^41^ This high sensitivity also renders the spectrometer well-suited for detecting other non-perturbative vibrational probes in this range such as N=N^+^=N^-^, or even the weaker C–D or C≡C stretches, which typically exhibit oscillator strengths up to tenfold lower than those of nitriles (Figs. S10-S12).^18, 21^ For example, azide-labeled GPCRs^62^ are an excellent candidate for *in cellulo* IR spectroscopy using the strategy described here and a QCL spectrometer for preservation of a native membrane environment.

Utilizing this spectrometer, we employed photoactive yellow protein (PYP) and its covalently attached chromophore p-coumaric acid (pCA) as a representative framework for many cysteine-based covalent inhibitors. For example, notable cysteine-reactive covalent drugs such as ibrutinib and acalabrutinib have transformed cancer therapy by selectively inhibiting Bruton’s tyrosine kinase (BTK) through a covalent mechanism, with minimal off-target effects on other kinases. ^63,64^ The PYP-pCA system provides a structurally and mechanistically analogous framework for investigating the molecular underpinnings of such selective covalent binding. While this work demonstrates covalent binding, the nitrile group’s sensitivity to its local environment enables detection of a broad range of molecular interactions, including non-covalent effects and drug molecules reliant on non-covalent binding. This versatility underscores a key advantage of this vibrational probe system over traditional fluorescence-based assays. Unlike fluorescence approaches that often require the attachment of bulky dye molecules—comparable in size to the drug itself—vibrational probes provide efficient, minimally invasive access to intrinsic molecular information, ranging from electrostatics to protein motion and conformational dynamics.

Such detailed structural insight is largely unique to spectroscopic techniques. In contrast, proteome-wide mass spectrometry assays, although powerful, face limitations. Highly specific biophysical studies of noncovalent interactions such as electrostatics or hydrogen bonding are not currently possible with mass spectrometry. Moreover, in this particular case, proteomic analysis aimed at identifying off-target binding interactions of pCA proved of limited utility. This was likely due to the standard reducing conditions for peptide linearization (involving iodoacetamide and dithiothreitol (DTT)), which can disrupt sensitive covalent linkages such as the thioester bond between PYP and pCA (SI Section S6).^65^ By comparison, the IR assay allows us to include those interactions typically challenging to mass spectrometry^66^—both non-covalent interactions plus covalent bonds sometimes susceptible to cleavage by DTT or tris (2-carboxyethyl)phosphine hydrochloride (TCEP) reducing agents employed in a standard proteomics digestion. This highlights the unique advantages of IR spectroscopy for probing the highly localized interactions that could potentially govern the chemical reactivity and efficacy of covalent drugs.

Finally, we performed an in-depth analysis of the unique spectral features of the nitriles observed using our QCL-based spectrometer and the PYP-pCA model system. Specifically, we detected distinct hypsochromic shifts in the nitrile stretching frequencies of variants F92oCNF and F96oCNF upon incorporation of the chromophore. To elucidate the origin of these shifts, we conducted molecular dynamics simulations using both fixed-charge and polarizable force fields, which revealed that the frequency changes stem from localized structural and electrostatic rearrangements within the protein environment upon pCA incorporation. These subtle perturbations highlight how the incorporation of a drug-like molecule such as pCA can induce site-specific changes in the protein’s electrostatic landscape—even far from the binding site—underscoring the sensitivity of vibrational probes to chemically relevant microenvironments. This becomes especially relevant when the local electrostatic environment of a binding site has been shown to be critical to the reactivity of covalent binding—a concept often overlooked by traditional electrostatic complementarity approaches used in drug design.^67^ Structurally, the crystallographic data seem to suggest a slight reorientation of F96 toward the empty chromophore pocket (Fig. 2) and slight increase in hydrogen bonding population (Fig. 4). In the case of F92oCNF, chromophore incorporation appears to increase its potential to form a hydrogen bond with T90, whereas in the *apo* state, it remains largely isolated from such interactions (Fig. 4). These results are consistent with an “induced fit” model of protein rigidification upon ligand binding, corresponding to an increase in internal hydrogen bonding networks near the ligand binding site—a trend potentially generalizable to a wide variety of drug targets.^68,69^

These findings validate the assay’s capacity to detect subtle structural perturbations resulting from covalent bond formation at reactive cysteine residues—a common motif in many modern covalent drugs. While our current proof-of-concept studies were conducted in bacterial cells, it is worth noting that bacterial proteins account for a significant fraction of therapeutic targets of interest in modern drug discovery.^70^ Furthermore, with emerging techniques in amber suppression, there is a growing potential to expand this assay to mammalian systems as well, where non-canonical amino acids can be incorporated efficiently.^71,72,73^ We further note the advantage in covalent drug uptake by mammalian cells, which do not have cell walls, as opposed to the *E. coli* we work with here—along with the relative stability of most true covalent drugs to hydrolysis compared to the challenges of pCA which must be chemically activated immediately prior to use.

Additionally, while in this study the vibrational probes were installed on the protein target, a similar assay could make use of vibrational reporters located on the drug itself— where a large portion of commercially available drugs already contain nitriles, alkynes, azides, or similar vibrational probes.^74^ Another foreseeable direction with this assay involves directed evolution to optimize molecular interactions in cells. For instance, we found that subculturing pCA-incubated cells in fresh media after the assay allowed for the regeneration of a new batch of cells available for further mutagenesis or future purification. Sensitive QCL spectroscopy enables the observation of vibrational frequency shifts in live cells, offering new opportunities to study molecular interactions in their native environments. These abilities can prove especially useful for understanding functions and properties of membrane and intrinsically disordered proteins, and more applications in which the native cellular environment is key.

## Supporting information

SI (Sections S1-S8) contains data on QCL spectrometer as-sembly, cell growth, protein purification, x-ray crystallog-raphy, FTIR spectroscopy, pCA act

## ASSOCIATED CONTENT

### Supporting Information

The Supporting Information is available free of charge on the ACS Publications website.

The SI (Sections S1-S8) contains data on QCL spectrometer assembly, cell growth, protein purification, x-ray crystallography, FTIR spectroscopy, pCA activation, mass spectrometry and proteomic analysis and computational data.

Spectrometer control MATLAB scripts are available for download at https://github.com/sdefried/QCL_spectrometer_scripts. Data analysis scripts for molecular dynamics simulations are available for download at https://github.com/KozuchLab/Publications/tree/main/oCNPhe_GROMACS_TINKER.

## AUTHOR INFORMATION

### Notes

Stanford University, which S.D.E.F., S.M., N.Y.H., N.B., and S.G.B. are affiliated with, has filed a patent application based on this work with S.D.E.F, S.M and S.G.B as inventors.

## ACKNOWLEDGMENTS

The authors thank Dr. Christopher K. Akhgar, Dr. Alicja Dabrowska, Dr. Andreas Schwaighofer, and Prof. Bernhard Lendl (TU Wien) for discussions on QCL spectrometer assembly. This work was supported in part by NIH Grant R35GM118044 (to S.G.B). S.D.E.F. is supported by the NSF Graduate Research Fellowship (DGE-1656518) and the Stanford CMAD Fellowship; N.B. by the Stanford Bio-X Bowes Fellowship; and J.K. by the DFG Individual Research Grant KO 5464-4 (project ID 493270578). Use of the Stanford Synchrotron Radiation Lightsource (SSRL/SLAC) was supported by the DOE Office of Science under Contract DE-AC02-76SF00515 and NIH Grant P30GM133894. Computational work was supported by the Sherlock cluster (Stanford) and ZEDAT HPC (Freie Universität Berlin, 10.17169/refubium-26754). We also thank the Stanford Chemistry NMR and SUMS facilities for pCA characterization. Special thanks to Mehmet Solyali (Stanford Machine Shop), Prof. Wei Min, Dr. Bing Xu, and Dr. Garvey McKenzie for their assistance with instrumentation, fruitful discussions and proteomics experiments, respectively.

## Footnotes

* We have shown that a combination of frequency shift and absolute intensity measurements can be used to obtain electric fields for nitriles;^23^ however, absolute intensity quantification is unreliable and difficult to perform in living cells.

