## Supplementary material for "Covalent Drug Binding in Live Cells Monitored by Mid-IR Quantum Cascade Laser Spectroscopy: Photoactive Yellow Protein as a Model System": SI (Sections S1-S8) contains data on QCL spectrometer as-sembly, cell growth, protein purification, x-ray crystallog-raphy, FTIR spectroscopy, pCA act

### Contents

**S1. QCL Spectrometer Assembly** (Figs. S1-S12, Tables S1-S4) 2
**S2. Cell Growth and Protein Purification** (Table S5) 15
**S3. X-ray Crystallography** (Fig. S13-S14, Table S6) 21
**S4. FTIR Spectroscopy** (Figs. S15-S18, Tables S7-S12) 25
**S5. Activation of p-Coumaric Acid** (Figs. S19-S25) 31
**S6. Mass Spectrometry** (Fig. S26) 37
**S7. Computational Details** (Fig. S27-S28) 39
**S8. Physical interpretability of nitrile shifts** (Fig. S29) 42
**Supplemental References** 43

### Section S1: QCL Spectrometer Assembly

**a. Laser performance and optical assembly:** The assembled double-beam transmission infrared spectrometer (Main Text Fig. 1C) uses an external-cavity quantum cascade laser (EC-QCL) as the light source (Daylight Solutions MIRcat-QT-2000 with M2048-P tunable module, cooled to 19°C using a closed-loop water chiller). The instrument design is based on the design of Akhgar et al.,[1] with several additional modifications that allow us to perform the desired analyses as described in the following sections. Firstly, we employ a different laser module to account for a new mid-IR spectral window (2000-2300 $cm^{-1}$, Fig. S1) in which the instrument is equipped to make high-sensitivity measurements with uniquely long pathlengths up to 250 µm when making measurements in water, where a weak but discernible IR absorption band is centered at roughly 2100 $cm^{-1}$. This is attributed to the combination of the HOH bending mode (~1630 $cm^{-1}$) with a librational mode (~400–700 $cm^{-1}$). The resulting combination band appears near 2100–2130 $cm^{-1}$. This unique capability of using longer pathlengths due to the high flux of this QCL module makes it ideal for analytes bearing oscillators such as C≡N, C–D, $N{=}N^{+}{=}N^{-}$, and C≡C stretches. This module provides pulsed output up to 30% duty cycle and high output power for high SNR: >1 W (peak) and >0.5 W (average). The laser control was implemented using custom scripts and MATLAB libraries provided by Daylight Solutions, which were modified for our specific use (Available at https://github.com/sdefried/QCL_spectrometer_scripts). For optical alignment, a collinear HeNe laser (632.8 nm) was used, which is integrated with the module and shares the same exit aperture as the infrared beam.

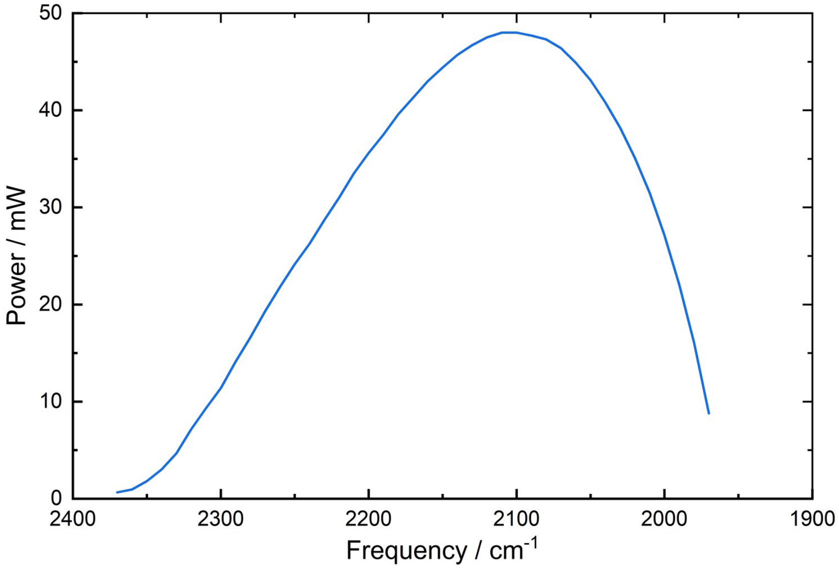


**Figure S1. Spectral power of the MIRcat-QT-2000 with M2048-P at 5% duty cycle:** Spectral power distribution of the MIRcat-QT-2000 with M2048-P tunable module, measured with a ThorLabs power meter measured at the exit aperture at 5% duty cycle and full current (780 mA). In the experiments to follow, a 30% duty cycle was used.

The laser beam enters a box containing all optics that is purged with dry air (Altec Air CO2-PG28 Purge Gas Generator operated at 60 psi). The beam is split by a 50:50 ThorLabs $CaF_2$ Plate Beamsplitter (Coating: 2–8 µm, Cat. No. BSW510). Mid-IR-enhanced gold mirrors (ThorLabs

Cat. No. PF10-03-M02) are used to define the optical path. Infrared-compatible neutral density filters (Andover Optics Cat. No. 150FNIR-25, 100FNIR-25, and 030FNIR-12.5) are used when applicable to equalize the intensity of the two beams or bring the overall power intensity into the dynamic range of the instrument (only when aqueous samples are not present that attenuate the light due to the libration and bending combination mode of water centered at ~2100cm$^{-1}$). $CaF_2$ diffusers were optionally included (Edmund Optics Cat. No. 19-734) to scramble polarization prior to detection. The two beams pass through sample and reference demountable liquid IR cells (Pike Technologies, Madison, WI, USA) designed for standard FTIR spectrometers (2 x 3" plate), which allowed a quick comparison of the same sample across different IR spectrometers used in the study. This cell features a needle plate with Luer-Lok fittings for convenient syringe filling with ~75 µL sample. The window dimensions are 32 x 3 mm, providing a 13 mm clear aperture. An O-ring seal model was utilized, employing two small O-rings around the window filling holes instead of a flat gasket. A pathlength of 250 µm for each cell was used by combining two circular Teflon spacers of 200 µm and 50 µm each. Cell holders are machined aluminum blocks that hold the demountable liquid cells in place using screws.

Inclusion of ND-filters and diffusers for balancing, and new $CaF_2$ focusing optics to increase incident power at the detector elements increase signal-to-noise specific for the detection of molecular vibrations in the spectral region (2000–2300 cm$^{-1}$). Note that the full power of the beam was used in conjunction with the 250-µm pathlength aqueous samples, with only attenuation of one beam relative to the other to equalize incident power on the detector. While the laser beam waist is < 2.5 mm, the beam is not focused prior to transmission through the sample cells, meaning the beam width is similar to but slightly less than that of the FTIR (6-mm aperture). The only focusing optics, consisting of $CaF_2$ plano-convex lenses (25.4 mm diameter, 100 mm focal length, Newport Optics Cat. No. CAPX13), are used to focus both the sample and reference beams after sample transmission into a custom-built thermoelectrically cooled balanced HgCdTe (MCT) detector, supplied by VIGO Photonics (PVI-4TE series), to detect the transmitted IR signal from both beams.[1] We note that further modification of the setup by using additional focusing optics may be possible for additional optimization.

Additional performance improvements may also arise from the specific detector modules used, as there is currently no single, unified head-to-head performance comparison published—either by manufacturers or third parties—between the VIGO PVI-4TE series and other commonly used liquid nitrogen-cooled MCT detectors, such as the Teledyne Judson models. However, detailed technical datasheets and catalogs are available for both to provide comparative performance metrics for HgCdTe detectors which we have summarized below. These resources allow for an indirect but robust comparison of key parameters such as detectivity, responsivity, noise, and spectral range.

**Table S1.** Parameters comparing thermoelectrically cooled VIGO HgCdTe detector used in QCL spectrometer and liquid nitrogen-cooled Teledyne Judson HgCdTe detector used in Bruker FTIR.

| Parameter | PVI-4TE (VIGO) | Teledyne Judson MCT (J15D) |
|---|---|---|
| **Cooling** | 4-stage TE | TE, $LN_2$, Stirling |
| **Spectral Range (µm)** | 2–12 (model-dependent) | 2–26 (model-dependent) |
| **Detectivity D* ($cm \cdot Hz^{1/2}/W$)** | ≥5×10⁹ (At $\lambda_{peak}$, TE; For 4TE-8 – used here) | Up to ~$10^{11}$ ($LN_2$, at $\lambda_{peak}$) |
| **Time Constant (ns)** | ≤3 (at $\lambda_{peak}$) | ~10–100 (model-dependent) |
| **Window** | Wedged ZnSe AR | Multiple options |
| **Area** | 1×1 mm² (typical) | 0.1–4 mm² (typical) |
| **Dark Current** | Very low (deep TE) | Low (TE), very low ($LN_2$) |

**b. Data acquisition, dynamic range and power calibrations:** The detector has three output channels—namely, signal (S), reference (R), and balanced (B = S – R) channels. Channel readout voltages from S, R and B are digitized by a high-speed lock-in amplifier (Stanford Research Systems SR865A 4 MHz dual-phase lock-in amplifier) whose reference signal is dictated by the laser pulse rate (1 MHz at 30% duty cycle). The lock-in amplifier sends capture buffers of S, R, and B covering laser scans from 2360-1970 cm$^{-1}$ to a host computer for analysis with an in-house MATLAB script that controls both data capture and laser operation. Digitized signal packets (Fig. S2) are processed with phase correction from the lock-in detector, Savitsky-Golay filtering, removal of anomalous spectra using similarity indices, scan averaging, and Fourier filtering, like that reported by Lendl and coworkers.[2]

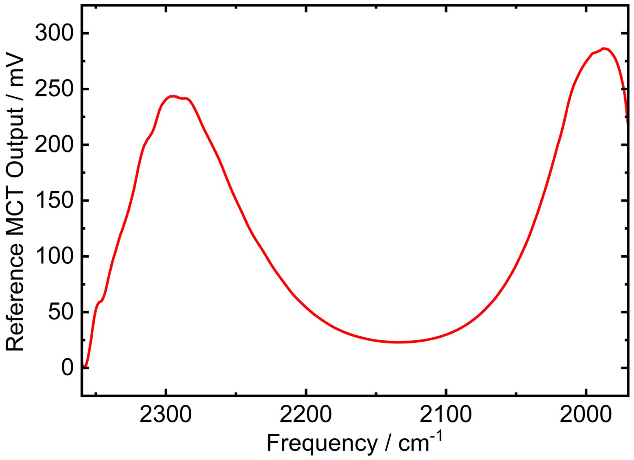


**Figure S2. Processed MCT output from QCL Module M2048-P scan through water-filled cell at 30% duty cycle:** Digitally processed reference MCT voltage output from 1970-2360 cm$^{-1}$ scan of the M2048-P tunable module at 30% duty cycle through a 250-µm sample cell filled with water. Although the laser's peak power is near 2100 cm$^{-1}$ (Fig. S1), water's combination mode at the same frequency significantly attenuates the beam in this frequency region. Output voltage was consistently < 300 mV for all spectral measurements reported herein.

Additionally, to increase sensitivity and make use of the full flux of photons from the laser (and extend the dynamic range of the detector), we externally calibrated the nonlinear detector voltage response in terms of laser power using a ThorLabs power meter (PM100D with S302C Thermal Power Sensor for 0.19 - 25 μm). During calibration, samples are absent in the beam path; therefore, up to three Andover Optics neutral density filters (optical densities 0.3, 1.0, and 1.5) were employed to attenuate the beam and maintain the MCT detector response within the linear dynamic range. At three different wavelengths, laser power was controlled by varying the current supplied to the QCL chip between 400 and 780 mA, while the beam intensity at both the signal and reference MCT were measured using both the MCT detector itself plus the thermal power meter. The detector voltage response to power demonstrates saturating behavior around 470 mV for both MCTs, which was well-captured using a rational instrument response function of the form $y = ax/(1 - bx)$ for $y$ = incident power at MCT and $x$ = MCT output voltage. The best-fit function to the data was used to convert measured detector voltage to beam incident power (Fig. S3). We note that this is also an addition beyond the capabilities of the instrument provided by Akghar et al.,[1] aimed to provide maximum possible photon flux and pathlength usage within our spectral range.

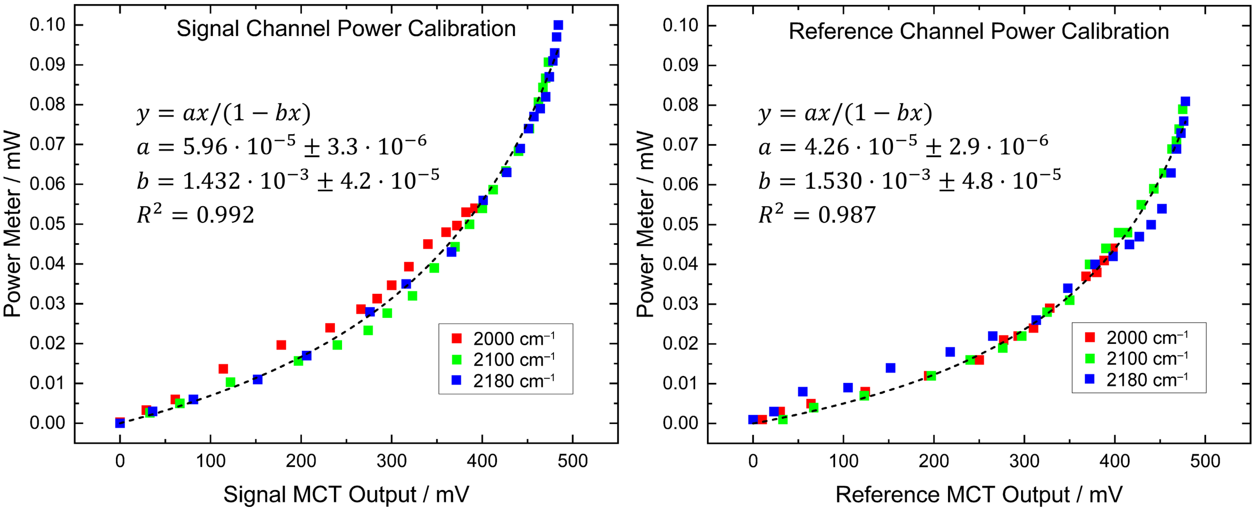


**Figure S3. Calibration of MCT detector output to incident power using nonlinear curve fitting:** Calibration against a thermal power meter is used to convert MCT detector output voltages to incident power on the detector, used to extend the dynamic range of the instrument given the nonlinear response of the detector at high photon incidence. Detector output voltages were consistently kept below 300 mV when transmitting through a 250-μm liquid sample cell filled with water, thus avoiding the saturation regime of the detector response, and increasing the accuracy of the transmission measurements. Fitting to the nonlinear response function was accomplished using the MATLAB curve fitting toolbox with a nonlinear least squares regression, fitting to all the data points (three separate wavelengths) at once. Errors reported are 95% confidence intervals of the parameters.

**c. Frequency calibration and data capture:** The frequency axes of the spectra were initially determined using the programmed laser start frequency (2360 cm$^{-1}$), at which a trigger was sent to the SR865A lock-in amplifier to begin a 32-kb data capture buffer, along with the SR865A capture rate (9765.6 Hz) and laser scan rate (2000 cm$^{-1}$/s). However, due to slight offsets between these frequencies calculated using time and the true laser frequencies, we recalibrated the frequency axis using a linear function. To do this, we used laser wavelength TTL triggers every 10 $cm^{-1}$ to subtract off from the MCT output as a separate channel in the lock-in amplifier, generating defined features in the voltage output capture buffer every 10 $cm^{-1}$. Correlating these with the original frequencies, we generated a linear calibration function that converted the time-based frequencies to corrected frequencies used as the axis for all absorption spectra reported herein (Fig. S4). By matching frequency signatures such benzonitrile to known literature values,[3] we verify that the calibration is accurate.

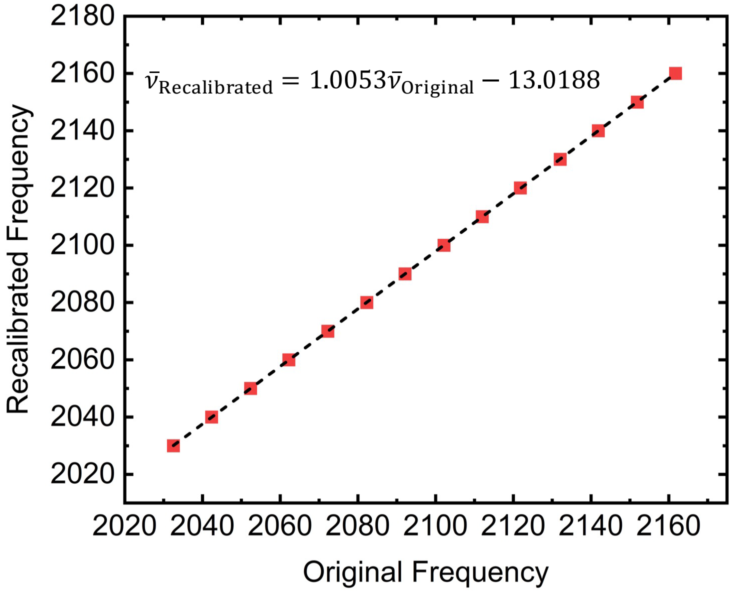


**Figure S4. Frequency calibration:** Recalibration of original laser scan frequencies determined by time from initial scan trigger, based on periodic wavelength triggers from the QCL.

**d. Absorbance measurements using the balanced detection scheme:** Absorbance of a sample relative to a reference blank was determined using a balanced detection scheme, which involves multiple measurements across different channels. Both a reference channel measurement of power versus laser frequency, and a balanced channel measurement of power difference versus laser frequency were obtained when reference blanks were loaded into both liquid cells. These are followed by loading the analyte of interest into the sample liquid cell and making a second balanced channel measurement of power difference versus frequency. Absorbance (A) is then calculated using the equation

$$\mathrm{A} = \log\left[P_{\text{Signal}}(V_{\text{Ref}} + V_{\text{Bal-Blank}}) / P_{\text{Signal}}(V_{\text{Ref}} + V_{\text{Bal-Sample}})\right],$$

where $V_{\text{Ref}}$ is the reference channel MCT output voltage and $V_{\text{Bal}}$ represents the balanced channel (subtracted) MCT output voltage, whether for both liquid cells filled with blanks ($V_{\text{Bal-Blank}}$) or the sample cell filled with sample against the reference filled with blank ($V_{\text{Bal-Sample}}$). The function $P_{\text{Signal}}(V)$ takes an MCT detector voltage as input and gives the beam power incident on the signal detector, according to the above calibration, $P_{\text{Signal}} = aV/(1 - bV)$. This method ensures that background noise is eliminated using balanced detection and that the measured absorbance is consistent with the Beer-Lambert law.

**e. Determination of experimental limit of detection:** Following the above process, we demonstrated the capabilities of the QCL spectrometer for sensitively measuring vibrational spectra of benzonitrile in water with a 250 μm pathlength, as shown in Fig. 1 of the main text. Spectra of benzonitrile at higher concentrations, measured on both the FTIR and QCL spectrometer, are shown below, along with the concentration/absorbance correlation (Fig. S5).

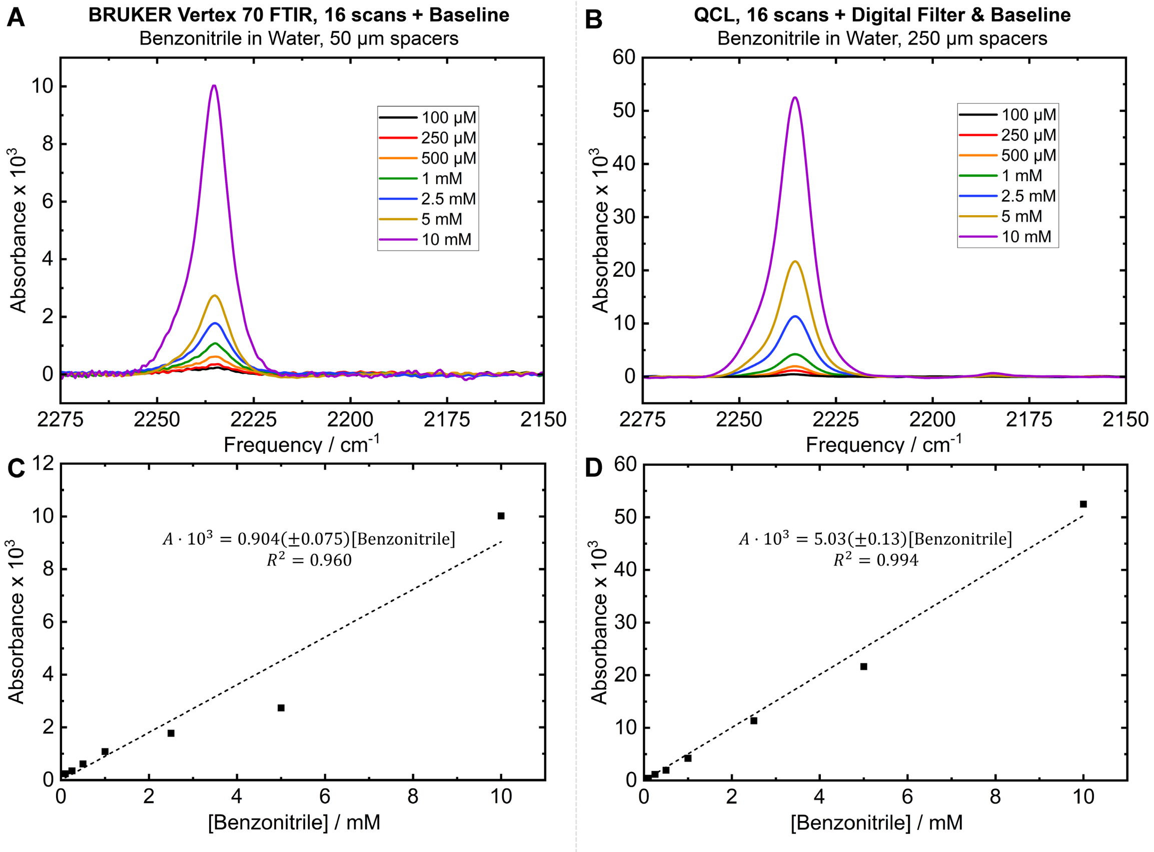


**Figure S5. Benzonitrile spectra and concentration–absorbance correlations from FTIR and QCL measurements:** Spectra of benzonitrile at concentrations of 100 μM – 10 mM are shown using 16 scans each on either (A) the Bruker Vertex 70 FTIR or (B) the QCL spectrometer. Correlation plots between concentration and absorbance are also shown for both (C) the FTIR and (D) the QCL spectrometer, with linear behavior according to the Beer-Lambert law, particularly in the case of the QCL spectrometer. Errors reported are standard errors. Note that the data for the FTIR carries a higher linear regression error compared to the QCL although the same samples were used for both. We attribute this to the relative difficulty in distinguishing peaks from solvent background when the absorption intensities are lower in the case of the FTIR.

The extinction coefficient of benzonitrile could be determined from both instruments. Firstly, the exact pathlength of both the FTIR and QCL spectrometer sample cells were determined using periodic spacings of etalon fringes as reported by Weaver et al.[3] The true pathlength differed slightly from the spacer widths (relating to how much the spacers were compressed), with the FTIR sample cell having pathlength 55.9 μm and the QCL sample cell having pathlength 259.4 μm. Using these values and the slopes reported below, extinction coefficients the benzonitrile nitrile stretch were calculated as $\varepsilon$ = 162 (±26) $M^{-1}$ $cm^{-1}$ for the FTIR and $\varepsilon$ = 194 (±10) $M^{-1}$ $cm^{-1}$ for the QCL (errors as 95% confidence intervals). We note that both values are within error of the $\varepsilon$ = 186 $M^{-1}$ $cm^{-1}$ value reported previously,[3] while meanwhile the peak maximum of 2235.5 $cm^{-1}$ (the more important metric for the current study) similarly agrees well, within 1 $cm^{-1}$, of our previous value for the frequency of benzonitrile in water.[3]

**f. Determination of theoretical limit of detection [LOD] and RMS noise analysis:** Using the calibrated instrument response to benzonitrile and the RMS noise from a blank spectrum (water versus water), we determined the theoretical limit of detection for benzonitrile for both the FTIR and QCL (Fig. S6, Table S2). The RMS noise ($\sigma_{\mathrm{RMS}}$) was measured over the range of 2260-2160 $cm^{-1}$, a relatively stable 100 $cm^{-1}$ window with high photon flux even through 250 μm aqueous samples, which also includes benzonitrile's peak of interest (Fig. S2). Theoretical limit of detection was defined by the concentration of benzonitrile required to achieve a signal at least three times the RMS noise over this region, that is, $[\mathrm{LOD}] = 3\sigma_{\mathrm{RMS}}/(\varepsilon \cdot l)$, for extinction coefficient $\varepsilon$ and pathlength $l$, where $\varepsilon \cdot l$ can be obtained from the slope of the instrument response function from Fig S5. As an additional point of comparison, we also include the scenario of detection based on the calibrated signal channel alone, without the use of balanced detection between two MCTs.

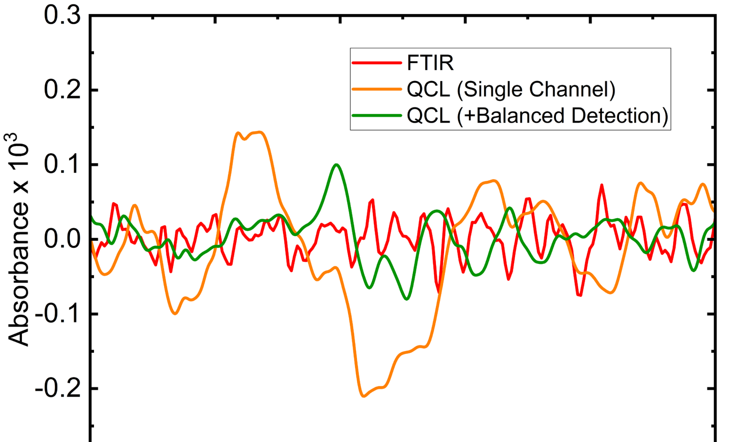


**Figure S6. Baseline spectra for estimating theoretical LOD.** Baseline IR spectra recorded by the QCL spectrometer using 16 scans, compared to the FTIR. Note that in all cases, 16 scans were used for ease of comparison, and the same procedure for digital processing of the QCL data was applied as described above. A manual baseline correction was employed to all spectra by fitting to broad, low-frequency polynomials to center the higher-frequency noise at zero across the spectral range.

**Table S2.** Limits of detection across different detection schemes (FTIR vs. QCL in balanced and single channel absorbance).

| | FTIR | QCL, Single Channel Only | QCL, Balanced Detection |
|---|---|---|---|
| $\sigma_{RMS}$ / Abs. Units | $2.4 \cdot 10^{-5}$ | $8.4 \cdot 10^{-5}$ | $3.0 \cdot 10^{-5}$ |
| **Benzonitrile LOD / µM** | 80 | 50 | 18 |

Our theoretical LODs determined from the baseline spectra (water minus water) and sensitivities match well with experimental spectra of benzonitrile at low concentrations (Fig S5 and Main text Fig. 1), where the nitrile stretch first becomes clearly visible to the FTIR between 50 and 100 µM, and to the QCL between 10 and 25 µM. The longer pathlength afforded by the higher photon flux is the primary advantage. As demonstrated above from the single channel results, the QCL light source inherently has more fluctuations than the FTIR globar source, an aspect observed by Lendl and coworkers in previous research,[1,2] although our use of a MIRcat laser equipped with ZeroPoint beam pointing control may provide some noise reduction compared to Lendl and coworkers' use of the Hedgehog laser. The ZeroPoint technology in the MIRcat provides active stabilization compared to the passive stabilization in the Hedgehog module and has lower thermal drift compared to the Hedgehog module.[4] In the absence of balanced detection, the Lendl Lab reported nearly a 3-fold higher RMS noise for their Hedgehog (2$^{nd}$ Gen) QCL source compared to FTIR systems under comparable experimental conditions, with measured values of $6.2\times10^{-5}$ (QCL single channel measurement, 53-s integration, 31-µm path length) versus $2.3\times10^{-5}$ (FTIR, 45-s integration, 8-µm path length) in the 1700–1500 cm$^{-1}$ spectral range. These results align closely with the noise levels reported in Table 1 of this work, despite differences in experimental parameters such as the higher number of scans (>100), shorter path lengths (8–31 µm) and a different spectral window employed in their study. The similarity in noise performance underscores the challenges inherent to QCL-based systems, where intensity fluctuations and thermal drift can dominate measurement uncertainty.

Despite the higher noise inherent to the QCL source, balanced detection decreases RMS noise into the realm of the FTIR, where the 5x longer pathlength of the QCL affords an approximately 5-fold advantage in terms of LOD. We note that in all cases described herein, 16 scans were used on the QCL spectrometer for comparison to the FTIR and measurement of the nitrile-embedded PYP within bacterial cells. This number was chosen as the optimal balance between noise-reduction and efficiency when making measurements (Fig. S7). Although Akhgar et al. demonstrated that 1000 scans could decrease noise by about 5x further beyond what we report here,[1] we aimed instead to minimize heating to our live cell samples by taking fewer scans. We found that 16 scans were sufficient to observe the nitrile peak within the cellular environments in each case.

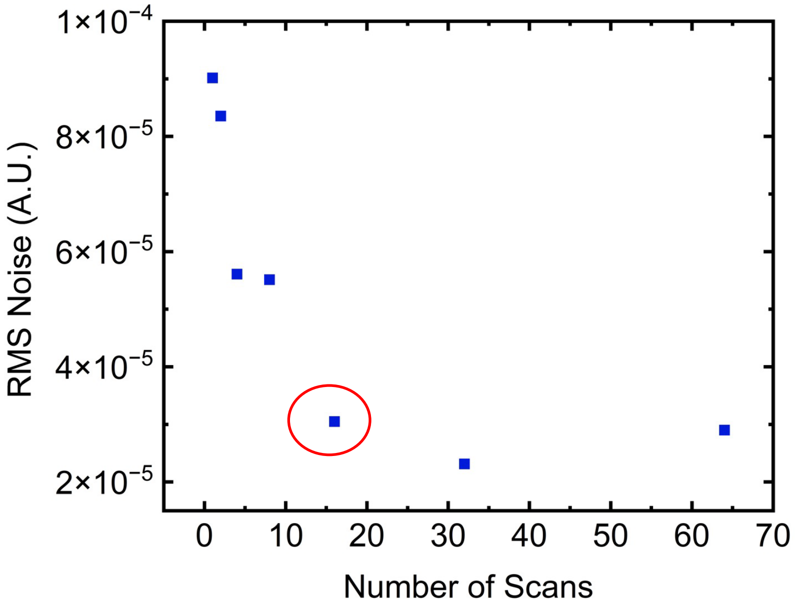


**Figure S7.** RMS noise for a blank (balanced) absorption spectrum through 250-μm pathlength water generally decreased with increasing number of scans based on square root law of noise averaging. We chose 16 scans as the optimal number of scans for the sake of achieving sufficient sensitivity while maintaining efficiency and minimal perturbation of the later cell-based samples.

In addition to the fewer scans, we also originally sought to reduce heating effects within the long-pathlength liquid sample cell by adding an extra 2-second dark time between scans, in addition to the typical 1.7-second time for scanning and transfer of the data capture buffer to the computer. We found no significant benefit to the additional equilibration time (Fig. S8, Table S3), indicating that on the timescale of these measurements, heating by the laser is not a major consideration. We note that further optimization of the setup should be possible to achieve a faster scan-time, such as via data streaming rather than use of a capture buffer or scanning over shorter spectral ranges (or fewer data points), in which case additional temperature controlling may be advantageous for the faster laser repetition rates. Currently, with a 1.7-second combined time for scanning over 300 $cm^{-1}$ and data transfer, this method proves roughly one-half as fast as a scan on a Bruker FTIR (0.8 seconds)—though still with markedly better signal-to-noise, meaning the QCL spectrometer still provides an overall benefit.

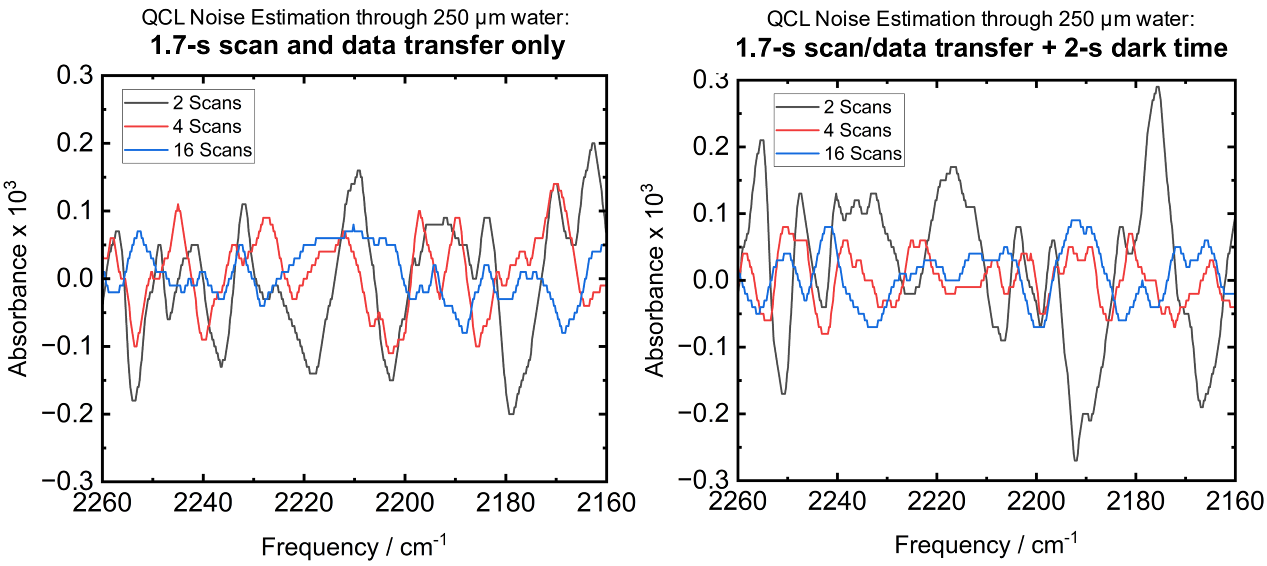


**Figure S8. Baseline spectra to compare RMS noise with additional dark time.** Baseline IR spectra recorded by the QCL spectrometer using 2, 4, or 16 scans, using (**a**) either the maximum repetition rate for the data buffer transfer (1.7 seconds), or (**b**) an additional 2 seconds (~3.7 seconds total) per scan for additional temperature equilibration. The same procedure for data processing and RMS noise calculation was followed as in Fig. S6, showing no major advantage from the additional equilibration time (and thus suggesting no heating artifacts using the 1.7-s laser repetition rate).

**Table S3.** RMS noise of baseline between 2260 and 2160 cm$^{-1}$, with varying numbers of scans and a variable dark time between scans for re-equilibration. Note that the variation between the two methods of detection does not appear significant, including compared to the previous reported baseline samples performed with 16 scans and the extra 2-s dark time (Table S1).

| **Number of Scans** | **1.7-s Scan/Data Transfer, $\sigma_{RMS}$ / Abs. Units** | **1.7-s Scan/Data Transfer + 2-s Dark-Time $\sigma_{RMS}$ / Abs. Units** |
|---|---|---|
| **2** | $4.1 \cdot 10^{-5}$ | $5.3 \cdot 10^{-5}$ |
| **4** | $2.6 \cdot 10^{-5}$ | $1.7 \cdot 10^{-5}$ |
| **16** | $1.7 \cdot 10^{-5}$ | $1.9 \cdot 10^{-5}$ |

As an additional point of comparison for our Bruker Vertex 70 FTIR, we also compared the detection of benzonitrile on an independent FTIR instrument, a ThermoFisher Nicolet iS50 with a liquid nitrogen cooled MCT detector. The sensitivity of this particular instrument, particularly in the benzonitrile region, was poorer than the Bruker Vertex 70 used for other comparative measurements reported in the study. Experimentally, the benzonitrile peak did not begin to be clearly visible in the spectrum until a concentration of ~500 µM, giving an experimental LOD several times higher than the 80 µM LOD on our Bruker FTIR (Fig. S9). We also calculated the RMS error for this instrument in the 2220–2245 cm$^{-1}$ region of the benzonitrile peak to be $4.4 \cdot 10^{-4}$ based on the baseline noise, an order of magnitude poorer than the Bruker FTIR. Additionally, the Nicolet iS50 requires ~1.6 seconds for each scan, twice as slow as the Bruker instrument. Hence, we conclude that the Bruker instrument's performance is a better comparison point for the QCL spectrometer performance. Note that this is an instrument-specific measurement for research benchmark purposes and not an endorsement of Bruker instruments.

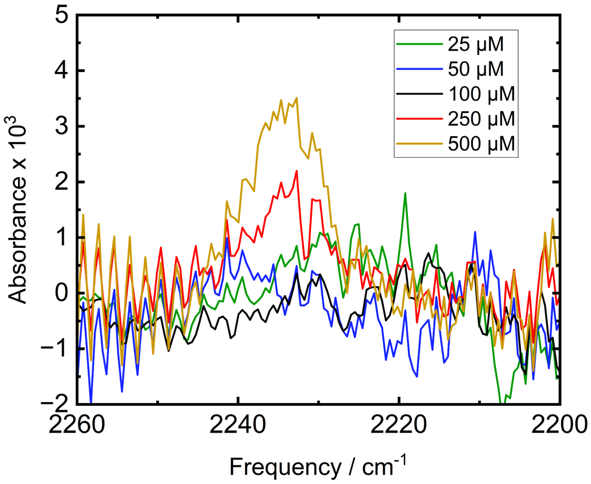


**Figure S9. Benzonitrile detection on a ThermoFisher Nicolet iS50 FTIR instrument.** Compared to the Bruker Vertex 70 FTIR, the ThermoFisher instrument requires a higher concentration of benzonitrile for detection and has a higher baseline noise, indicating that the Bruker Vertex 70 FTIR is a better comparison point for the QCL spectrometer performance.

**g. Extending to alkyne C≡C, C–D, and $N{=}N^{+}{=}N^{-}$ stretches:** In addition to nitriles, several other vibrational stretches that can function as readouts of molecular interactions and environments and that have been introduced into proteins by amber suppression are also reasonable candidates for detection using this QCL spectrometer at low concentrations. The 2000–2300 $cm^{-1}$ region is home to other weak vibrational oscillators including carbon-deuterium (C–D) alkyne (C≡C) and stronger azide ($N{=}N^{+}{=}N^{-}$) bonds that have proven useful probes in a variety of biological contexts.[5,6] We demonstrate how our instrument can sensitively detect sample vibrational probes at other neighboring frequencies including an alkyne (1-hexyne) at 2111 $cm^{-1}$, a deuterated (aldehyde-H) form of N-cyclohexylformamide (CXF-D) at 2182 $cm^{-1}$ used as a probe of electric field directionality in enzymes by Zheng et al.,[7] and an azide-labeled phenylalanine. As can be seen in Fig. S10, the QCL measurement of 10 mM 1-hexyne in water using a 250 µm pathlength gives a significantly better spectrum compared to the same number of scans (16) on the FTIR using a 50 µm pathlength. We moreover show how the QCL can give good sensitivity for detection of strongly absorbing azides such as 4-azidophenylalanine (Fig. S11). Finally, we show how the QCL again gives better sensitivity for detection of the CXF-D, with a clearly visible peak at 1 mM where the same number of scans on the FTIR at the same concentration and 50 µm pathlength does not yield a similarly clear peak (Fig. S12). Comparisons of the peak fitting parameters using either method are provided in Table S4.

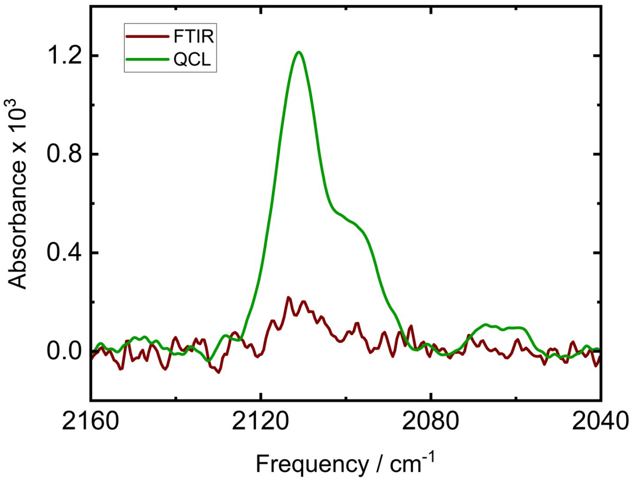


**Figure S10.** Spectra of 10 mM 1-Hexyne in water after 16 scans on either the Bruker Vertex 70 FTIR using a 50 µm pathlength, or on the QCL using a 250 µm pathlength, show better sensitivity of the QCL in this frequency range.

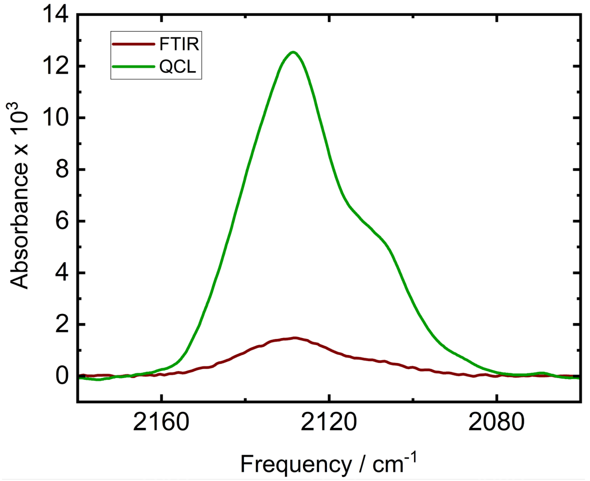


**Figure S11.** Spectra of 1 mM 4-azidophenylalanine in water after 16 scans on either the Bruker Vertex 70 FTIR (here using 50 and 25 µm spacers to reduce etalon fringes), or on the QCL using a 250 µm pathlength, show better sensitivity of the QCL in this frequency range.

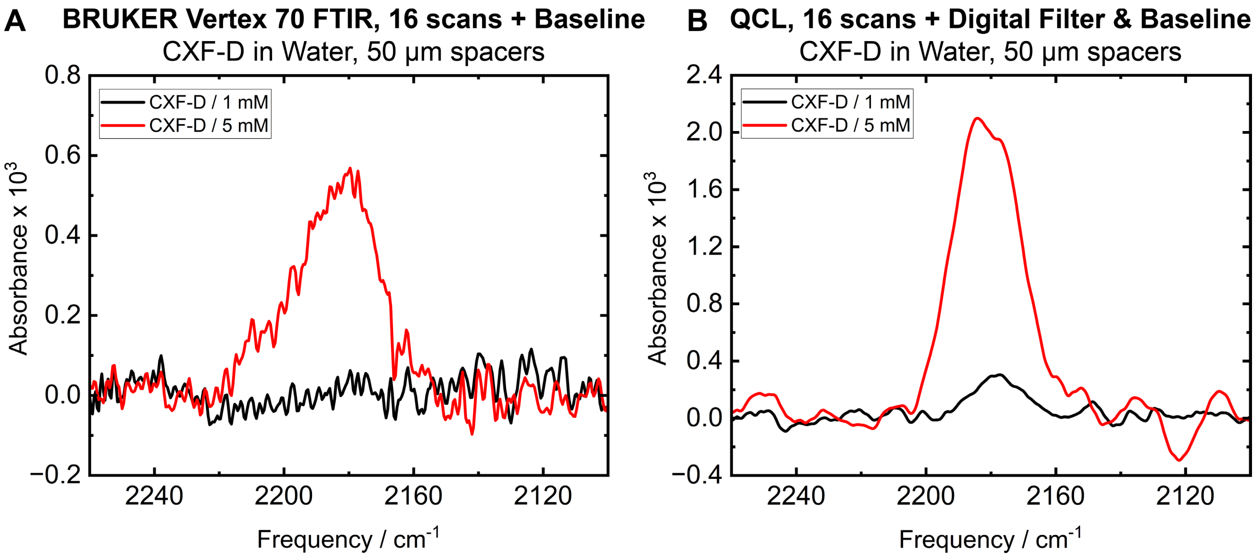


**Figure S12**. Spectra of CXF-D at concentrations of 1 mM and 5 mM in water to compare sensitivity of (A) the Bruker Vertex 70 FTIR at a 50 µm pathlength, and (B) the QCL spectrometer at a 250 µm, with 16 scans used on both instruments.

**Table S4.** Peak fitting parameters from Bruker Vertex 70 FTIR and QCL (balanced detection) measurements. Data in this table are reported for analyte concentrations where peaks were clearly identifiable in both techniques, with 16 scans performed in each case to ensure comparability. Peaks are fit in OPUS using Gaussian + Lorentzian functions of variable weights (Voigt profiles).

| **Analyte** | **Bond** | **Concentration** | **FTIR** | | | **QCL (Balanced)** | | |
|---|---|---|---|---|---|---|---|---|
| | | | Peak / $cm^{-1}$ | Width / $cm^{-1}$ | Intensity/ OD | Peak / $cm^{-1}$ | Width / $cm^{-1}$ | Intensity / OD |
| **Benzonitrile** | **C≡N** | 100 $\mu$M | 2237.7 | 16.8 | $2.1 \cdot 10^{-4}$ | 2235.8 | 8.6 | $4.4 \cdot 10^{-4}$ |
| **1-Hexyne** | **C≡C** | 10 mM | 2111.1 | 11.2 | $1.8 \cdot 10^{-4}$ | 2111.4 | 12.1 | $1.2 \cdot 10^{-3}$ |
| | | | 2097.6 | 8.1 | $7.6 \cdot 10^{-5}$ | 2097.2 | 12.2 | $4.4 \cdot 10^{-4}$ |
| **CXF-D** | **C–D** | 5 mM | 2183.6 | 29.9 | $5.3 \cdot 10^{-4}$ | 2181.3 | 24.1 | $2.1 \cdot 10^{-3}$ |
| **4-Azido-phenylalnine** | **N=N=N** | 1 mM* | 2129.6 | 26.3 | $1.5 \cdot 10^{-3}$ | 2129.7 | 25.9 | $1.2 \cdot 10^{-2}$ |
| | | | 2105.3 | 17.6 | $3.3 \cdot 10^{-4}$ | 2105.8 | 17.1 | $3.8 \cdot 10^{-3}$ |

*FTIR measurements for 4-Azidophenylalnine used 50 µm and 25 µm spacers to reduce fringes, all others used 50 µm spacers for FTIR and 250 µm for QCL.

**Section S2: Cell growth and protein purification**

**a. Sequence of PYP:** N-terminal 6X-His-tagged PYP with an enterokinase cleavage site, from a previously reported pBAD construct,[3] was recombinantly expressed in *E. coli* both as WT and in various nitrile-incorporated variants using Amber suppression. The highlighted positions were individually replaced with the TAG stop codon for Amber suppression machinery. The positions code for Phe residues at F28, F62, F92 and F96.

ATGGGGGGTTCTCATCATCATCATCATCATGGTATGGCTAGCATGACTGGTGGACAGC
AAATGGGTCGGGATCTGTACGACGATGACGATAAGGATCCGAGACTCGAGGATGACG
ATGACAAAATGGAACACGTAGCCTTCGGTAGCGAGGACATCGAGAACACCCTCGCCA
AGATGGACGACGGCCAGCTCGACGGCCTGGCCTTCGGCGCCATCCAGCTCGACGGC
GACGGCAACATCCTTCAGTACAACGCCGCGGAGGGCGACATCACCGGCCGCGACCC
GAAGCAGGTCATCGGCAAGAACTTCTTCAAGGACGTGGCCCCGTGCACTGACAGCC
CGGAGTTCTACGGCAAGTTCAAGGAAGGGGTGGCCTCGGGCAACCTGAACACGATG
TTCGAGTACACCTTCGATTACCAAATGACGCCCACGAAGGTGAAGGTGCACATGAAG
AAGGCCCTCTCCGGCGACAGCTACTGGGTCTTCGTCAAGCGCGTCTAA

**b. Mutagenesis and cloning:** MAX Efficiency™ DH10B competent cells (ThermoFisher) underwent a double transformation. This involved introducing two plasmids: pEVOLpylT-N346AC348A and one of several pBAD-PYP variants with/without the engineered stop codons for nitrile incorporation (Addgene). The PYP plasmids, including those with the TAG mutations, were created using an Agilent QuikChange Lightning Site-Directed Mutagenesis Kit, starting from the original pBAD-PYP plasmid. A separate mutation, C69A (for chromophore knockout), was generated using a Q5 Site-Directed Mutagenesis Kit from New England Biolabs. The primers used for making the C69A mutations were the following: C69A Forwards: GCGACTGACAGCCCGGAGTTC and C69A Reverse: CGGGGCCACGTCCTTG. Annealing temperature for this reaction was set to 68ºC.

**c. Cell growth without Amber suppression**: Plasmids for both wild-type PYP and the PYP-C69A mutant (without Amber suppression machinery) were transformed into competent Bl21 (DE3) *E. coli*. Transformed cells were selected by overnight growth on agar plates containing 100 µg/mL ampicillin at 37°C. Single colonies were used to initiate 5 mL liquid cultures in LB with 100 µg/mL ampicillin, which were grown overnight at 37°C with shaking at 250 rpm. These overnight cultures were then used to inoculate 1L cultures in either Luria Bertani Broth or Terrific Broth (ThermoFisher), supplemented with 100 µg/mL ampicillin, in pre-warmed baffle flasks (Nalgene). These larger cultures were grown at 37°C with shaking at 200 rpm until they reached an OD600 of 0.6 (typically 2-4 hours). Protein expression was then induced by adding 143 mg of IPTG (0.6 mmol, ThermoFisher), and the cultures were incubated overnight at 20°C with shaking at 200 rpm.

**d. Cell growth using Amber Suppression**: The general protocol for sample preparation involved the incorporation of noncanonical amino acids, ortho-cyanophenylalanine (oCNF, ChemImpex, Ann Arbor, MI, USA), into overexpressed target proteins within *E. coli* utilizing an Amber suppression system. Doubly transformed cells, harboring plasmids for both the target protein and the orthogonal translation machinery, were initially cultured following established literature procedures.[3] For each liter of subsequent protein expression, a starter culture was prepared by inoculating a 14 mL round-bottom culture tube containing 5 mL of Luria-Bertani (LB) growth medium (Millipore) with the transformed cells. These starter cultures were grown for 12 to 16 hours, or until they reached saturation, with the LB medium supplemented with specific antibiotics for plasmid selection: 100 μg/mL ampicillin for the pBAD plasmid and 34 μg/mL chloramphenicol for the pEVOL plasmid. Following sufficient growth, the entire starter culture was transferred into 1 L of auto-induction medium from which phenylalanine source had been omitted. This larger volume of auto-induction medium was also supplemented with the same antibiotic concentrations: 100 μg/mL ampicillin (Sigma-Aldrich) and 34 μg/mL chloramphenicol (Sigma-Aldrich) and pre-warmed to 37°C for one hour prior to inoculation. It is important to note that the auto-induction medium contained arabinose (Sigma-Aldrich), which was added after the medium had been sterilized via autoclaving to prevent degradation of the arabinose necessary for the pBad plasmid expression. The inoculated cultures were then shaken at 250 RPM at 37°C for 30 minutes, after which 10 mL of a 100 mM solution of oCNF was added. 100 mM oCNF solution was prepared approximately 1 to 2 hours before its addition to the cell culture medium using sonication and heating as oCNF is only partially soluble in water at these concentrations. Protein expression was then auto induced for approximately 19 hours at 250 RPM and 37°C.

**e. Preparation of cells for QCL-based spectroscopy**: After overnight expression of the cultures, a portion of ~100 mL liquid culture was introduced to freshly prepared activated pCA (acyl imidazolide form, see section S5), scraping the yellow oil product from the flask walls and mixing with the liquid cell culture, before stirring overnight at 4°C.[3] The control (non-exposed) cells were similarly incubated overnight but without pCA. The following morning, both groups of cells were washed 3x (gentle centrifugation at 2000 RPM for 10-15 minutes, followed by resuspension) with minimal media (Sigma Aldrich M9 Minimal Salts; Cat. No. M6030) to remove excess oCNF and/or pCA. In the pCA-incubated batch, a noticeable yellow coloration of the loose cell pellets collected in each washing step provided a visual indication of the successful incorporation of the chromophore into the *apo*-PYP protein within the cells. Notably, excess activated pCA, which did not react with *apo*-PYP, underwent hydrolysis in water present in the medium, regenerating insoluble pCA. This re-formed pCA presented itself as a distinct white precipitate layer, often observed above or as a separate layer from the denser yellow cell pellet after the initial centrifugation (confirmed using IR, NMR and UV-vis). To isolate the cells containing the bound chromophore, the layer of the pellet containing the yellow cells was carefully scraped off using a spatula, thereby separating it from the white precipitate, for subsequent washing with minimal media in parallel with the non-pCA-incorporated cells.

Following the washing steps, the loose cell pellets were resuspended in a minimal amount of M9 minimal media. Aliquots were diluted 1000x for measurement of $OD_{600}$ by a PerkinElmer Lambda-365 UV-visible spectrophotometer (Waltham, MA), a common metric to assess the concentration of a bacterial suspension. The final $OD_{600}$ of the washed bacterial suspensions when diluted 1000x were adjusted to the range of $OD_{600}$=0.3-0.4 for measurements. Approximately 75 µL of the $OD_{600}$-adjusted suspension was loaded into the sample demountable cell of the QCL spectrometer using a syringe. Cellular suspensions expressing the wild-type protein were prepared and washed in a similar manner for reference measurements. Using the previously described protocol for the QCL measurements, IR spectra were collected for the F28oCNF, F62oCNF, F92oCNF, and F96oCNF variant-expressing cells that had been incubated with or without the activated pCA, taken with reference to wild-type *apo*-PYP-expressing cells (lacking the nitrile reporter) serving as a crucial reference blank to identify the nitrile vibrational bands reporting on pCA binding.

**f. Acquisition of nitrile IR spectra in live cells**: Averaged spectra from 16 scans for each of the different nitrile-containing mutants were collected to assess the microenvironment around a ligand of interest. In all cases, the laser pulse rate was kept at 1 MHz with 300 ns duration (30% duty cycle), with scanning from 2360 $cm^{-1}$ to 1970 $cm^{-1}$ at a scan rate of 2000 $cm^{-1}/s$. A 2 second dark time was employed between consecutive scans to allow for data transfer and temperature re-equilibration (minimal over the time duration of each scan, See Figure S8). Data was collected in 16-kb data packets on the SRS865A lock-in amplifier at a rate of 9765.6 Hz (equal to the maximum collection rate divided by 16). The time constant of the SRS865A lock-in amplifier was kept as 300µs, the input range 1 V, and the sensitivity 500 mV. Data packets were sent via ethernet streaming to the host computer for processing. Data processing was based on the methods reported by Lendl and coworkers,[2] and included optional phase correction in cases where the signal was significantly distributed between *x*- and *y*- components (typically not necessary), plus removal of anomalous spectra using similarity indices (rarely employed in practice), a 3rd-order Savitsky-Golay filter on 41 data points at a time, averaging of successive scans, and a sharp (order-400) long-pass Fourier Filter with a Blackman-Harris window and 1000 Hz cutoff over the spectral window.

**g. Protein purification:**

- ***Apo*-PYP for vibrational spectroscopy:** To purify *apo*-PYP for measurements, the following protocol was employed: Cell pellets obtained from centrifugation were resuspended in a buffered 20 mM Tris-HCl, 10 mM NaCl (Buffer A; pH 8.0) using 2 mL of buffer per 1 mL of cell pellet. Cells were lysed by homogenization using a C3 Emulsiflex, and insoluble cell debris was removed by high-speed centrifugation at 15,000 g for 120 minutes. The resulting lysate was filtered through a 0.22 µm Sterile PVDF filter and then loaded onto a Ni-NTA column for 6x-His tag affinity purification. The column was equilibrated with 5 column volumes (CVs) of Buffer A (1 CV = 5 mL resin) before sample loading. The protein was allowed to bind to the column for 1 hour under gentle rocking. Subsequently, the column was washed with 5 CVs each of Buffer A containing 10 mM and 20 mM imidazole to remove non-specifically bound entities. The 6X-His-tagged PYP was then eluted with Buffer A containing 250 mM imidazole. To remove excess imidazole, the eluted protein was subjected to buffer exchange using spin filtration with Amicon Ultra 15 centrifugal filter units (3 kDa MWCO) against Buffer A. The protein concentration was determined by absorbance at 280 nm using the Expasy ProtParam-calculated extinction coefficient for the linker+his-tagged *apo*-PYP (MW=18501.5 g/mol; $\varepsilon_{280}$=14400 $M^{-1}$ $cm^{-1}$), and the purity assessed using LC-MS with a C8 reverse-phase column and single quadrupole mass spectrometer at the SUMS open access facility at Stanford University.

- ***Apo*-PYP for crystallography:** For x-ray crystallography studies, further purification steps were performed to remove the His-tag. The Ni-NTA purified *apo*-PYP was subjected to anion exchange chromatography (Akta FPLC, GE Healthcare) using an FPLC system with three 5 mL Q HP columns (Cytiva) connected in series, pre-equilibrated with Buffer A. Protein elution was achieved by applying a linear gradient of 0–40% Buffer B (1 M NaCl, 20 mM Tris-HCl, pH 8.0) over 180 mL at a flow rate of 5 mL/min. Fractions eluting between 25% and 32% Buffer B were pooled. These pooled fractions were concentrated and buffer-exchanged against Buffer A using 15 mL 3 kDa MWCO Centrifugal Filter Units (Millipore) to remove excess NaCl. The protein was then concentrated to ~10 mg/mL, and the N-terminal 6X-Histag was cleaved by adding 1 Unit of enterokinase (New England Biolabs, Catalog #P8070S) per 0.025 mg of protein and 2 µL of 1 M $CaCl_2$ per 1 mL of protein solution. The cleavage reaction was performed under gentle shaking at 25 RPM 4 °C for 16 hours. Following tag cleavage, the mixture was re-purified using anion exchange chromatography as described after the Ni-NTA step. Untagged PYP fractions eluted between 15% and 19% Buffer B. Fractions exhibiting maximal $A_{280}$ were pooled. As before, NaCl was removed by buffer exchange against Buffer A using a 3 kDa Amicon centrifugal filter unit. For samples intended for protein IR spectroscopy, *apo*-PYP was further concentrated using a 0.5 mL 10 kDa Amicon centrifugal filter unit by reducing the volume to approximately 18 $\mu$L and then centrifuging for an additional hour at 14,000 g at 4 °C. The concentrated *apo*-PYP was carefully collected, gently pipetted to ensure homogeneity, and transferred to a 0.6 mL microcentrifuge tube prior to crystallography discussed in Section S3.

- ***Holo*-PYP for vibrational spectroscopy:** To purify *holo*-PYP, two distinct approaches were employed for different confirmatory measurements within the study. In the first method, adapted from the original protocol by Weaver et al.,[3] the crude *apo*-PYP cell lysate obtained from one liter of expressed cells was directly reacted with the activated pCA chromophore immediately after the evaporation of THF. The activated chromophore, which adhered to the sides of the round-bottom flask in the form of a dense yellow oil, was scraped using a metal spatula into the cell lysate. This mixture was gently stirred (< 200 RPM) overnight to avoid introducing foam. The following morning, insoluble reaction precipitate, primarily a white solid form of excess pCA was removed by centrifugation at 15,000 g for 120 minutes. The resulting lysate was then filtered and purified using the Ni-NTA column as described previously for *apo*-PYP. In the second protocol for *holo*-PYP purification, cells where the activated pCA was intended for incorporation were first lysed, and then the purification procedure was repeated as described for *apo*-PYP using the Ni-NTA column.

**h. Estimation of sensitivity of in-cell IR measurements:** In their work on detection of proteins via the Amide-I and Amide-II bands, Akhgar et al.[1] report an effective limit of detection (LOD) of 0.0025 mg/mL for three proteins: hemoglobin, gamma-globulin, and concanavalin A. We convert their protein LODs to molar concentrations of amides based on their numbers and this is summarized in Table S5, which we use to compare to our benzonitrile LODs.

**Table S5.** Estimation of molar concentrations of amides from protein concentration LODs of Akhgar et al. We assume the extinction coefficient of amides ~1000 $M^{-1}cm^{-1}$.

| Protein | ~MW (kDa) | Residues | Protein Concentration based on 0.0025 mg ml$^{-1}$ (μM) | Approx. Amide Concentration [Protein Conc x Residues] (μM) |
|---|---|---|---|---|
| **Hemoglobin** | 65 | 578 | 0.038 | 22 |
| **IgG** | 150 | 1020 | 0.017 | 17 |
| **Concanavalin A** | 104 | 948 | 0.024 | 23 |

As shown above, at 0.0025 mg/mL protein, Akhgar et al. could detect amides at concentrations around ~20 μM. Our setup utilizes a 250 μm Teflon spacer, roughly ten times longer in path length than theirs (26 μm). While this longer path length is beneficial to us, nitriles have an oscillator strength approximately five times lower than that of amides (~200 vs. 1000 $M^{-1}$ $cm^{-1}$).[3] Consequently, if the numbers reported in the setup of Akhgar et al. were completely translatable to our case of nitriles, we would anticipate a benzonitrile LOD half that of carbonyls at the short pathlength, or ~10 μM. This value is similar in magnitude to the 18 μM LOD for benzonitrile we report here. There are significant differences between the two apparatuses, most significantly in the type of laser module used over the different wavelength ranges (our MIRcat being more powerful and stable compared to the Hedgehog), and the significantly lower number of scans we used in the current work (16 scans here versus 1000 scans reported by Akhgar et al., which would give them an approximately 8-fold advantage for reducing random noise due to scan replicates). While a more exact comparison is not possible, a theoretical LOD at a similar magnitude to that anticipated by the apparatus of Akhgar et al. helps confirm the theoretical limitations of balanced QCL-based spectrometers across a variety of chemical species and wavelengths.

**i. Estimation of protein concentration in cells:** From the purification of 1 L cultures, we obtained 10 mg of *apo*-F92oCNF and 16 mg of *apo*-F96oCNF (both 6X-His-tagged) PYP. Assuming similar growth conditions and expression systems, $OD_{600}$ measurements of the liquid cultures taken after 19 hours across three biological replicates (diluted 50x, with $OD_{600} = 1$ corresponding to $8 \times 10^8$ *E. coli* cells/mL) yielded average cell densities of $3.8 \pm 1.5 \times 10^9$ cells/mL for cells expressing *apo*-F92oCNF and $6.6 \pm 3.0 \times 10^9$ cells/mL for *apo*-F96oCNF. The protein per cell for both *apo*-F92oCNF and *apo*-F96oCNF was calculated based on the protein yield and average cell count, with uncertainties in the cell counts considered. For growths expressing *apo*-F92oCNF, the protein per cell was found to be approximately $2.6 \times 10^{-15} \pm 1.0 \times 10^{-15}$ g/cell, which corresponds to $1.4 \times 10^{-19} \pm 5.7 \times 10^{-20}$ mol/cell of the tagged *apo*-PYP. For *apo*-F96oCNF, the protein per cell was $2.4 \times 10^{-15} \pm 1.1 \times 10^{-15}$ g/cell, corresponding to $1.4 \times 10^{-19} \pm 6.2 \times 10^{-20}$ mol/cell. The relatively small difference in protein per cell, with overlapping error margins, suggests that the expression levels for *apo*-F92oCNF and *apo*-F96oCNF are similar in protein production per cell, corresponding to about 84000 protein molecules per cell. If we then assume each cell is ~1 $\mu m^3$, this results in a concentration of nitrile per cell at 140 μM, which is within the limit of detection of our QCL-spectrometer for individual cells. Recognizing that the cell suspension will be more dilute in terms of nitrile-labeled *apo*-PYP than an individual cell, we consider the concentration of cells within the final cell suspension based on a 1000x dilution giving $OD_{600}=0.35$. Again using $OD_{600} = 1$ corresponding to $8 \times 10^8$ *E. coli* cells/mL, this corresponds to $2.8 \times 10^{11}$ cells/mL in the cell suspension used for the QCL measurements. Using $1.4 \times 10^{-19}$ mol protein/cell, the final nitrile concentration visible to the QCL detection assay was thus calculated as 39 μM in each case—above the 18 μM detection limit for the QCL spectrometer but still smaller than the FTIR LOD for benzonitrile using the same number of scans. While we note that the extinction coefficient of the oCNF in the organized protein environment will not be identical to benzonitrile in solution, these results indicate an ability and advantage of the QCL in detecting vibrational frequencies of nitrile-tagged *apo*- or *holo*-PYP in live cell suspensions.

**j. Comparison to another protein system, GFP:** To show that our method is generally applicable even with proteins with different expression levels and in a different chemical environment, we considered another protein system with embedded nitriles, to show similarly that IR spectra could be recorded for the nitrile-labeled protein in a different cellular environment. Green fluorescent protein (GFP) has been a topic of significant interest—particularly in terms of properties governing its photophysics, with roles of electrostatics and sterics in tuning properties such as absorption and emission wavelength, lifetime, and fluorescence quantum yield.[8,9] Most FPs autocatalyze their chromophore maturation within a chromophore forming triad **XYG**. In previous work from our group, we successfully incorporated various non-canonical Y analogs into the GFP chromophore, including halogenated tyrosines, tyrosines with strongly electron-withdrawing $NO_2$ groups, electron-donating ortho-methoxy groups, and in this work with ortho-cyanophenylalanine (oCNF) giving a unique nitrile stretching frequency. The exact structural and photophysical implications of this observation are beyond the scope of the current work and will be pursued elsewhere. Olenginski et al. have recently reported the successful incorporation of para-cyanophenylalanine (pCNF) into the GFP chromophore.[10]

To create this construct, we introduced a stop codon (TAG) at position Y66 (pBAD-sfGFP-66TAG construct). MAX Efficiency™ DH10B Competent Cells (ThermoFisher) were co-transformed with the pEVOL-pylT-N346A/C348A plasmid (Addgene) and the pBAD plasmid for protein expression via Amber suppression. Defined media supplemented with ampicillin, chloramphenicol, arabinose, and oCNF (as a phenylalanine analog) were used for cell growth. These conditions mirrored those used to generate oCNF analogs of PYP. The cells were auto induced and grown for 18 hours at 37 °C, after which the temperature was reduced to 18 °C for 4-6 hours to facilitate chromophore maturation. Following expression, sample preparation and the measurement protocol followed similarly to the PYP experiments.

### Section S3: X-ray crystallography

**a. Comparison to previous *holo*-PYP crystal structures**: X-ray crystallography was used to compare the global structure of *apo*-PYP to the chromophore-bound state. The chromophore-bound state has been previously crystallized as wild-type at ultra-high resolution (PDB: 1NWZ, resolution = 0.82 Å)[11] and as the four various nitrile-containing variants we use here (resolution < 1.2 Å): F28oCNF (PDB: 7SPX), F62oCNF (PDB: 7SPW), F92oCNF (PDB: 7SPV), and F96oCNF (PDB: 7SJJ).[3] These four variants station the nitrile probe at various distances from the chromophore binding site and in both hydrogen-bonding and non-hydrogen-bonding environments, as shown in Fig. S13 below.

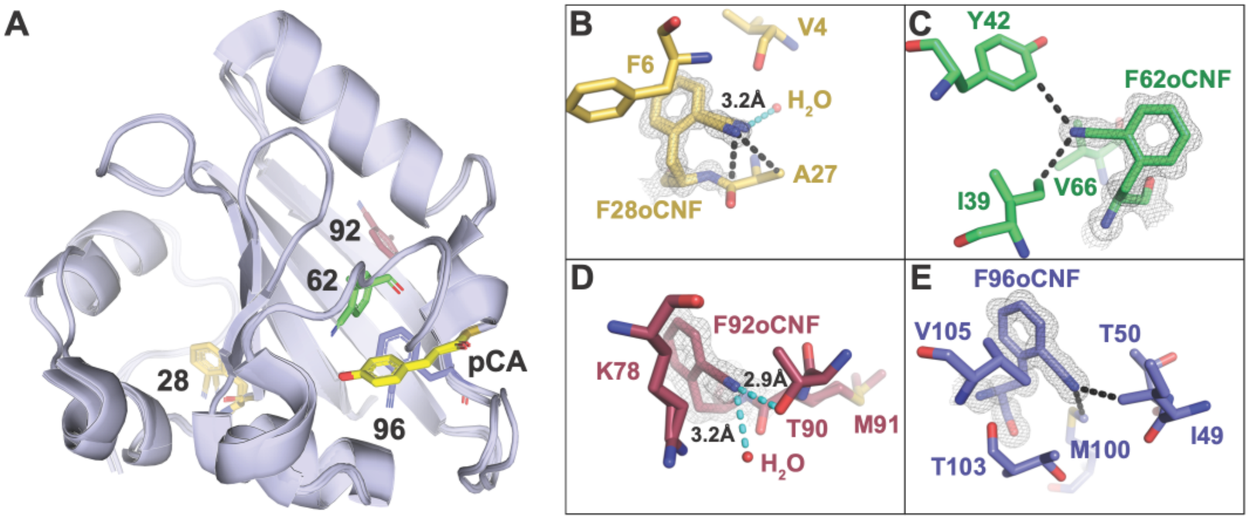


**Figure S13**. (A) Superimposed structures of the four nitrile-containing variants of *holo*-PYP, with the individual nitrile environments shown in detail: (B) F28oCNF interacts with one water molecule in its crystallographic pose, (C) F62oCNF lies in a hydrophobic environment, (D) F92oCNF demonstrates two hydrogen bonds to the nitrile in the crystallographic pose, from T90 and water, creating a strong electric field, and (E) F96oCNF demonstrates no hydrogen bonding partners in its crystallographic pose. Figure partially reproduced from Kirsh et al. (ACS Publications).[12]

**b. Methods of *apo*-PYP Crystallography, Diffraction, and Analysis:** In the present work, *apo*-PYP was crystallized using the hanging-drop method as previously described for *holo*-PYP.[13] Freshly purified *apo*-PYP was buffer-exchanged into a buffer of pH 6.0, 20 mM potassium phosphate (Sigma-Aldrich) and concentrated to 20 mg/mL. Hanging drops were set manually using 24-well VDX plates with sealant (Hampton Research) and 22 mm diameter siliconized glass coverslips (Hampton Research). 1 µL of protein was mixed with 1 µL of mother liquor on each coverslip and closed over wells containing 500 µL mother liquor. Trays were kept at room temperature and covered in aluminum foil to maintain consistency with *holo*-PYP crystallization conditions. Mother liquor solutions contained 1M NaCl and ammonium sulfate (Sigma-Aldrich) ranging from 1.8 – 2.6 M in 0.1 M increments. Crystal formation was observed within one to four weeks of initial setting.

All crystals of sufficient quality were placed on 0.1 – 0.4 mm Mounted CryoLoops (Hampton Research), briefly (1-2 seconds) incubated in polyfluoroether cryo-oil (Hampton Research), rapidly frozen in liquid nitrogen, and placed into a Stanford Synchrotron Radiation Lightsource (SSRL) cassette for X-ray diffraction data collection. X-ray diffraction was performed at SSRL (Menlo Park, CA) Beam Line 12-1 at 100K. Further detailed information on data collection strategy can be found in Romei et al.[13] Data processing and scaling was performed using the HKL3000 software, providing the correct C2 space group.[14] Molecular replacement was performed in Phenix-1.21.2 with phenix.phaser using the WT PYP (PDB Entry: 1NWZ) structure as the search model.[15] Numerous rounds of model building and refinement were carried out in Coot32[16] and then with phenix.refine. Refinement continued until the R-free score could no longer be lowered. The X-ray collection parameters and refinement statistics are given below (Table S6).

**Table S6.** Crystallographic data summary

| **Entry name** | *Apo*-PYP |
|---|---|
| **PDB Entry** | 9O8V |
| **Beamline** | SSRL 12-2 |
| **Wavelength (Å)** | 0.97946 |
| **Detector Distance (mm)** | 150 mm |
| **Space Group** | C 1 2 1 |
| **Unit Cell Dimensions** | 104.63 Å, 50.72 Å, 71.25 Å<br>90.00°, 132.93°,90.00° |
| **Total Observations** | 310844 (12896) |
| **Unique Reflections** | 66798 (3224) |
| **Redundancy** | 4.7 (4.0) |
| **Completeness (%)** | 98.5 (95.2) |
| **Mean I/σ** | 42.2 (1.55) |
| **Wilson B-factor (Å²)** | 17.7 |
| **$R_{meas}$** | 0.073 (0.811) |

| | |
|---|---|
| **$CC_{1/2}$** | 1.00 (0.729) |
| **Reflections used in Refinement** | 66760 |
| **Reflections used for $R_{free}$** | 3445 |
| **$R_{work}$** | 0.1580 |
| **$R_{free}$** | 0.1853 |
| **Number of non-H Atoms** | Protein – 1770, Ligands – 0, Solvent – 177 |
| **Protein Residues** | 117 |
| **RMSD Bond Lengths (Å)** | 0.013 |
| **RMSD Bond Angles (°)** | 1.352 |
| **Ramachandran Favored (%)** | 97.35 |
| **Ramachandran Outliers (%)** | 2.65 |
| **Rotamer Outliers (%)** | 0.00 |
| **Clashscore** | 3.51 |
| **Average B-factor, all atoms (Å$^2$)** | 25.0, Protein –24.0, Chloride – 35.9, Waters– 37.3 |

**c. Structural comparisons between *apo*- and *holo*-PYP:** The global structure of *apo*-PYP, as revealed by its crystal structure (PDB ID: 9O8V), shows minimal deviation from the *holo* form (PDB ID: 1NWZ), even in the absence of the pCA chromophore (or any incorporated nitriles). A comparative B-factor Z-score analysis between the *apo* and *holo* structures using the $C_\alpha$ carbons was performed to quantify local structural fluctuations. This analysis revealed generally similar patterns of disorder between the two forms. The B-factors here serve as a tool to assess differences in atomic fluctuations between two protein structures. B-factors (also called temperature factors) extracted from crystallographic data reflect atomic displacement or static disorder (a consequence of imperfect unit cell ordering in the crystals). Normalized B-factors were employed because different crystals often have different amounts of static disorder. To perform a Z-score analysis, the B-factors are normalized using the formula:

$$Z = \frac{B_i - \mu}{\sigma},$$

where $B_i$ is the B-factor of residue $i$, $\mu$ the mean B-factor across the structure, and σ is the standard deviation. The resulting Z-scores help identify regions with statistically significant differences in disorder. This analysis is presented in Figure S14, where we note that, excluding the N-terminal and C-terminal regions of the *apo* form (9O8V) that are either unresolved or appear disordered, most of the other residues in both the *holo* or *apo* form have comparable Z-scores.

To further characterize structural similarity, we performed fixed-charge molecular dynamics simulations for 100 ns using two different starting configurations: (1) the *holo*-PYP structure with the chromophore coordinates removed (PDB ID: 1NWZ), and (2) the native *apo* structure (PDB ID: 9O8V). These results are presented later in Figures S24-S25; comparative analysis of these trajectories revealed no substantial conformational shifts or secondary structure rearrangements, regardless of the initial structure used. The minimal structural deviations observed throughout both

MD trajectories indicate that global conformational changes upon pCA binding are negligible. These techniques are more structurally informative than previous circular dichroism (CD) studies on *apo* and *holo* PYP,[17] where the CD spectrum in the 250-300 nm region is dominated by contributions from the chromophore in the *holo* form, making structural comparison from the protein secondary structure difficult. These findings validate the computational strategy of modeling *apo*-PYP variants by using *holo*-PYP structures and simply removing the pCA chromophore coordinates. This approach was subsequently adopted in our AMOEBA polarizable force field simulations of *apo*-PYP variants, with detailed simulation protocols provided in Section S7.

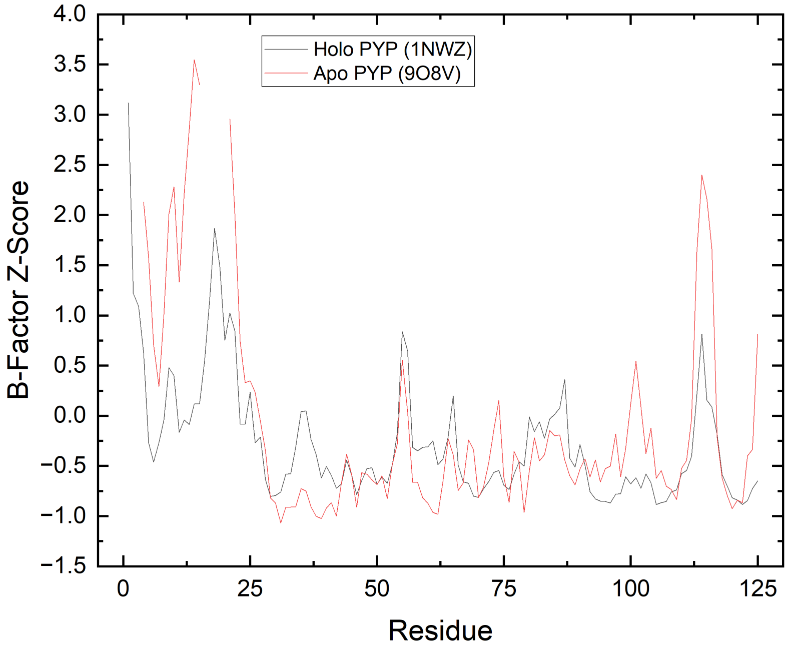


**Figure S14.** B-factor Z-score analysis comparing the crystal structures of *holo*-PYP (PDB ID: 1NWZ, chromophore incorporated, red) and *apo*-PYP (PDB ID: 9O8V, grey). The deviations are generally similar between the two forms. Notably, the N-terminal and C-terminal regions in the *apo* form are either missing or disordered. Overall, the crystal structures suggest minimal differences in local disorder, a conclusion further supported by the MD simulations presented in Section S7.

**Section S4: FTIR spectroscopy**

**a. Methods and materials:** For FTIR measurements involving small molecules (e.g., benzonitrile), the molecule was dissolved in solvent, where a stock solution of 10-100 mM was prepared and consequently diluted for LOD measurements. Both 1-hexyne and benzonitrile were sonicated for 10 minutes by an ultrasonication water bath to ensure complete dissolution.

For proteins (e.g. PYP), the protein was exchanged into a buffer containing 20 mM Tris HCl and 10 mM NaCl at pH 8.0. PYP was concentrated using 0.5 mL 10 kDa Amicon centrifugal filter units to approximately 18 µL, at which point it was further concentrated for an additional hour at 14,000 g at 4°C. Samples were transferred to Eppendorf tubes, and the volume was recorded. The final concentration of the protein was adjusted to ~10mM using this protocol. In either case (small molecule or protein), ~15-20 µL of the prepared sample was then injected into a demountable IR cell, which was assembled using two $CaF_2$ optical windows (19.05 mm diameter, 3 mm thickness, Lambda Research Optics, Inc.) separated by a pair of Teflon spacers (25 µm and 50 µm thick). FTIR spectra were recorded using a Bruker Vertex 70 spectrometer with a liquid nitrogen-cooled mercury cadmium telluride (MCT) detector (Teledyne Judson, Inc.), and the sample chamber was constantly purged with dry air to remove atmospheric water. The spectra were recorded in a wavenumber window spanning 4000–1000 $cm^{-1}$ with 512 scans at 1 $cm^{-1}$ spectral resolution. The obtained FTIR transmission spectra were converted to the corresponding absorption spectra using the neat solvent/background spectra taken under similar conditions as the reference. Data collection and processing were performed by the spectroscopy software OPUS 5.0.

**b. FTIR experiments with cells for *apo*- and *holo*-F96oCNF PYP:** We considered the ability of the FTIR to measure nitrile probe frequencies within the expressing cellular environment as we did with the QCL spectrometer. We estimated that the working concentration of nitrile-tagged proteins within the cell suspension was higher than the theoretical LOD (using that of benzonitrile, See Table S4), while it was less clear whether peaks could be observed at all using the FTIR with a shorter pathlength. As an aside, we observed that transmission in the FTIR through 250-µm pathlength sample cells filled with water was too low and below the noise floor in this region. Using the 50-µm pathlength, we were able to pick out the nitrile peak of both *apo*- and *holo*-F96oCNF PYP within the cell suspension from the background using 16 scans as the QCL; however, the spectra were significantly noisier and thus difficult to extract quantitative data in terms of frequency shifts between the *apo*-protein in cells and pCA-bound protein in cells (Fig. S15, Table S7). Qualitatively, a blue-shift does appear to occur upon pCA-binding, agreeing with the QCL-based results, though the lack of clarify in these spectra is a strong point in favor of using the QCL-afforded sensitivity for these measurements.

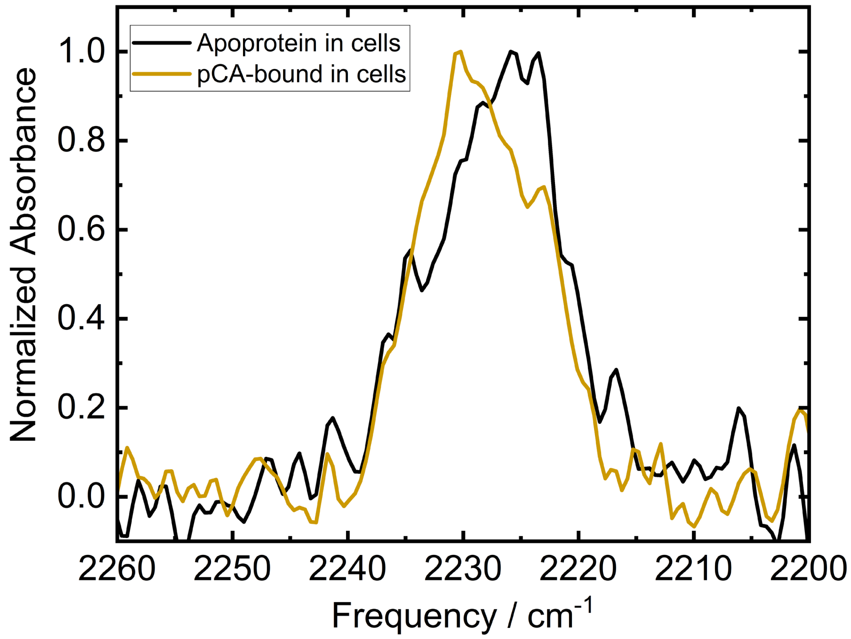


**Figure S15.** FTIR spectra of *apo*-F96oCNF PYP (black) and pCA-incubated F96oCNF PYP (gold) in expressing *E. coli* cells, showing the same qualitative blueshift upon pCA binding, albeit with significantly more noise.

**Table S7.** Spectral data fits, F96oCNF PYP in cells using FTIR. Peaks are fit in OPUS using Gaussian + Lorentzian functions of variable weights (Voigt profiles).

| Protein | Peak / cm$^{-1}$ | Width /cm$^{-1}$ | Intensity/OD |
|---|---|---|---|
| **F96oCNF *Apo* Cells** | 2226.9 | 14.2 | $3.1 \cdot 10^{-4}$ |
| **F96oCNF pCA Cells** | 2228.5 | 12.7 | $3.6 \cdot 10^{-4}$ |

**c. Free oCNF spectrum (FTIR) and comparison to *in cellulo* Y66oCNF sfGFP (QCL Spectra):** Using a similar measurement protocol as for the PYP samples, we recorded 16 QCL-based IR scans of oCNF-GFP-expressing cells versus unlabeled superfolder-GFP-expressing cells as a reference. The scans revealed a clear peak at 2225.8 cm$^{-1}$, corresponding to the nitrile incorporated into the chromophore—distinct from free oCNF (Fig. S16)—thereby establishing another cellular system amenable to direct probing with this spectrometer.

To confirm that the observed peak arose from the nitrile-tagged GFP rather than free oCNF, we compared the vibrational spectra of the cell suspensions and purified protein with that of free oCNF (50 mM aqueous solution), collected using the Bruker Vertex 70 FTIR (black trace). The 2232.1 cm$^{-1}$ peak of free oCNF is sufficiently distinct from the GFP spectra to confirm that the observed vibrational signature originates from the nitrile-labeled protein and not residual oCNF (which had been thoroughly washed from the cells). Further confirmation was obtained using purified proteins. The proteins were isolated from cells via Ni-affinity chromatography after 6X-His tag cleavage. Cells were pelleted at 15,000 × g for 1 hour following homogenization. The supernatant was filtered using a 0.22 µm filter and loaded onto 5 mL of Ni-NTA resin. After incubation, the proteins were eluted using 250 mM imidazole, as described for PYP in the preceding sections. The eluted proteins were then buffer exchanged using Amicon 10 kDa spin filters and further concentrated to 10 mM for FTIR analysis.

An FTIR spectrum of the purified protein (10 mM), acquired using a Bruker Vertex 70 spectrometer, displayed a single peak at 2224.9 cm$^{-1}$. The ~1 cm$^{-1}$ red shift relative to the in-cell measurements may indicate slight differences in the cellular context (e.g., pH or salt concentration) compared to the purified protein or differences in maturation of the chromophore due to kinetics or the conditions of purification. The enhanced sensitivity of the QCL spectrometer enabled these types of measurements.

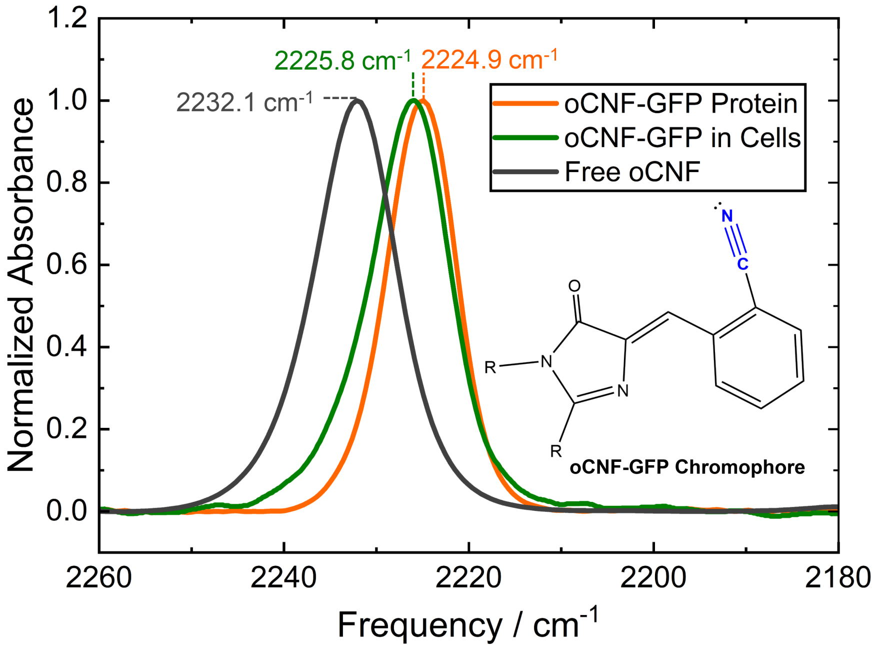


**Figure S16.** Comparison of QCL-based IR spectrum of Y66oCNF-superfolder GFP within expressing cells (green) to FTIR spectrum of 10 mM purified protein (orange) and of 50 mM free oCNF in water (black). Frequencies are analyzed according to peak maxima.

**d. FTIR spectroscopy of PYP variants:** In addition to the QCL-based measurements of the nitrile-tagged proteins in expressing cells, we also compare these results to FTIR measurements of the purified proteins, proteins in cells, and a relevant mutant. In general, the QCL-based spectra of pCA-bound *holo*-PYP proteins within the expressing cellular environments (Tables S8-S9) match well (consistently within 0.6 cm$^{-1}$) of the peak centers for the isolated *holo*-PYP proteins reported previously by Weaver et al.[3] We also verify the similarities between the QCL-based spectra of *in cellulo* proteins and the FTIR-based spectra of purified PYP proteins—whether *apo*-PYP or the pCA-incorporated PYP after incubation of the cells with PYP for the purposes of the QCL experiments (Fig. S17, Tables S10-S11). Notably, the *in cellulo* PYP spectra are also distinct from the 2232.1 cm$^{-1}$ for free oCNF (Fig. S16), verifying that these spectral features derive from nitrile-tagged PYP rather than free oCNF in aqueous solution. In general, the nitrile frequencies for PYP in the cell and the isolated protein are within 2 cm$^{-1}$ of one another, with the exception of *apo*-

F92oCNF where the *in cellulo* nitrile frequency is red-shifted by ~4 $cm^{-1}$. Though in both cases the *apo*-form is significantly red-shifted compared to the pCA-bound form, this difference appears to be larger (by ~ 4 $cm^{-1}$) in the cell than in the isolated protein. Because the F92oCNF results are highly dependent on a dynamic hydrogen-bonding equilibrium involving the nitrile and neighboring T90 as discussed in the main text and evidenced by AMOEBA simulations (Section S7), it is possible that this equilibrium could be highly dependent on not just the chromophore presence but also the environmental conditions such as pH, ionic strength, or crowding/osmotic stress environment. Though the results agree qualitatively whether inside cells or *in vitro*, the slight differences give rise to a view as to how *in cellulo* measurements may be better-poised to capturing the true conformational ensemble of the protein, before or after ligand binding, in live cells.

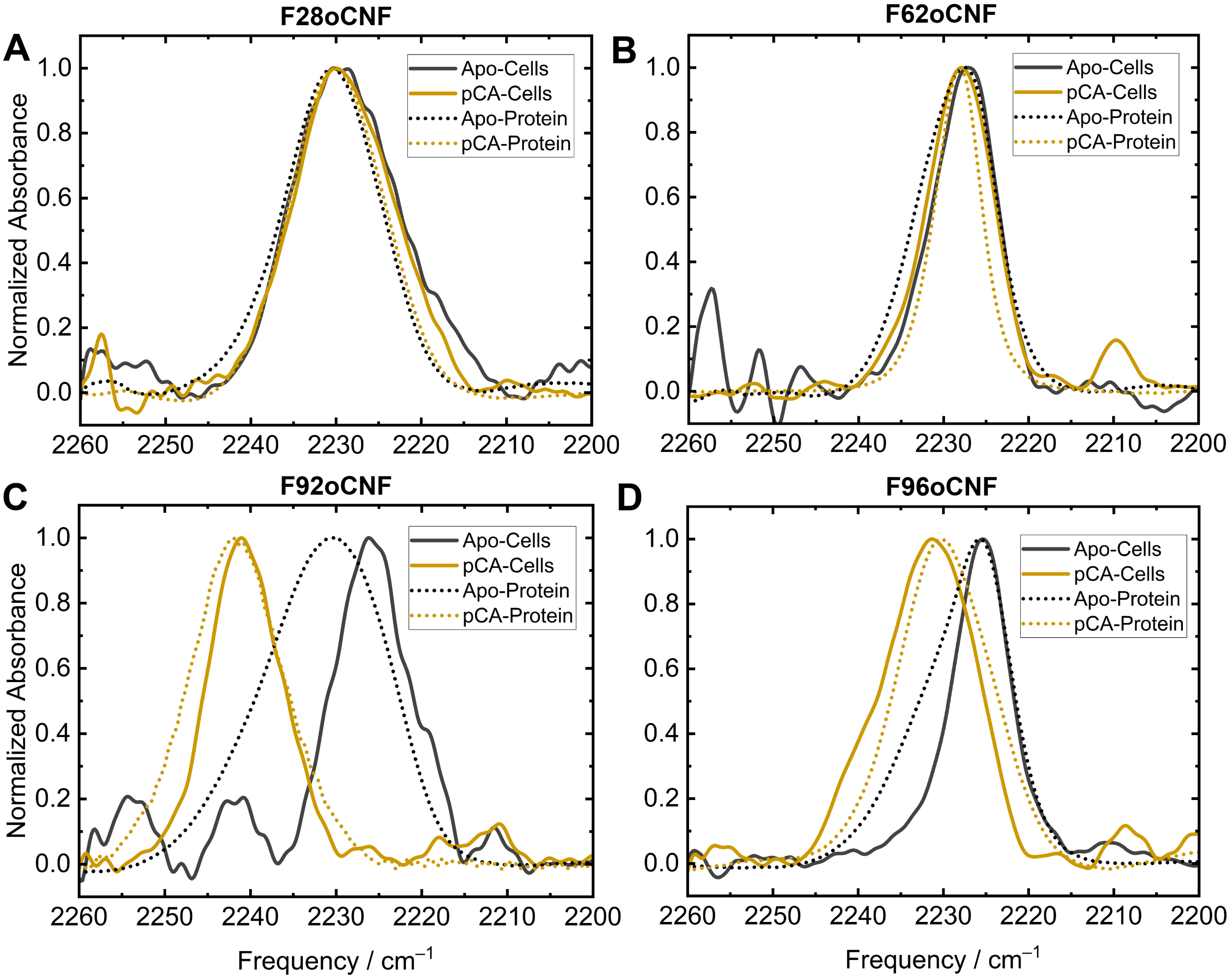


**Figure S17.** PYP spectra with or without incubation of cells with pCA—the *in cellulo* measurements from the QCL (solid line) are compared to FTIR measurements on the purified proteins (dotted lines). In each case, the purified proteins whose FTIR spectra are shown are the proteins extracted from the same cell batches used for the *apo* or pCA-incubated whole-cell spectroscopy, with the exception of F62oCNF, for which we show the FTIR spectrum from a previous purification by Weaver et al.[3]

**Table S8.** Observed frequency of the nitriles in PYP variants with and after pCA addition in cells using the QCL spectrometer. Frequencies are analyzed here according to **curve fitting.**

| PYP Variant | *Apo*-PYP $\bar{\nu}_{Nitrile}$ / cm$^{-1}$ | *Apo*-PYP Width / cm$^{-1}$ | pCA-PYP $\bar{\nu}_{Nitrile}$ / cm$^{-1}$ | *Apo*-PYP Width / cm$^{-1}$ | $\Delta\bar{\nu}$ After pCA |
|---|---|---|---|---|---|
| **F28oCNF** | 2228.8 | 14.3 | 2229.1 | 13.9 | **+0.3 cm$^{-1}$** |
| **F62oCNF** | 2227.5 | 8.7 | 2228.1 | 8.9 | **+0.6 cm$^{-1}$** |
| **F92oCNF** | 2225.8 | 9.7 | 2240.7 | 9.7 | **+14.9 cm$^{-1}$** |
| **F96oCNF** | 2225.4 | 6.8 | 2231.7 | 12.5 | **+6.3 cm$^{-1}$** |

**Table S9.** Observed frequency of the nitriles in PYP variants with and after pCA addition in cells using the QCL spectrometer. Frequencies are analyzed according to the **peak maxima**.

| PYP Variant | *Apo*-PYP $\bar{\nu}_{Nitrile}$ / cm$^{-1}$ | pCA-PYP $\bar{\nu}_{Nitrile}$ / cm$^{-1}$ | $\Delta\bar{\nu}$ After pCA |
|---|---|---|---|
| **F28oCNF** | 2230.5 | 2230.5 | **0 cm$^{-1}$** |
| **F62oCNF** | 2227.3 | 2227.9 | **+0.6 cm$^{-1}$** |
| **F92oCNF** | 2226.2 | 2241.1 | **+14.9 cm$^{-1}$** |
| **F96oCNF** | 2225.3 | 2231.3 | **+6.0 cm$^{-1}$** |

**Table S10.** Observed frequency of the nitriles in PYP variants with and after pCA addition after purification using 10mM protein on the FTIR spectrophotometer. Frequencies are analyzed according to **curve fitting.**

| PYP Variant | *Apo*-PYP $\bar{\nu}_{Nitrile}$ / cm$^{-1}$ | *Apo*-PYP Width / cm$^{-1}$ | pCA-PYP $\bar{\nu}_{Nitrile}$ / cm$^{-1}$ | *Apo*-PYP Width / cm$^{-1}$ | $\Delta\bar{\nu}$ After pCA |
|---|---|---|---|---|---|
| **F28oCNF** | 2230.5 | 13.1 | 2229.9 | 12.7 | **– 0.6 cm$^{-1}$** |
| **F62oCNF** | 2228.4 | 10.7 | 2228.3 | 6.5 | **– 0.1 cm$^{-1}$** |
| **F92oCNF** | 2231.1 | 16.8 | 2241.6 | 13.0 | **+10.5 cm$^{-1}$** |
| **F96oCNF** | 2227.1 | 12.1 | 2229.9 | 12.8 | **+2.8 cm$^{-1}$** |

**Table S11.** Observed frequency of the nitriles in PYP variants with and after pCA addition after purification using 10mM protein on the FTIR spectrometer. Frequencies are analyzed according to the **peak maxima**.

| PYP Variant | *Apo*-PYP $\bar{\nu}_{Nitrile}$ / cm$^{-1}$ | pCA-PYP $\bar{\nu}_{Nitrile}$ / cm$^{-1}$ | $\Delta\bar{\nu}$ After pCA |
|---|---|---|---|
| **F28oCNF** | 2230.4 | 2230.3 | **– 0.1 cm$^{-1}$** |
| **F62oCNF** | 2227.6 | 2228.2 | **+0.6 cm$^{-1}$** |
| **F92oCNF** | 2230.5 | 2242.3 | **+11.8 cm$^{-1}$** |
| **F96oCNF** | 2225.9 | 2230.2 | **+4.3 cm$^{-1}$** |

**e. FTIR experiments using a PYP C69A control variant:** In addition to FTIR controls involving the isolated nitrile-embedded proteins, we also verified that the *apo*-protein spectra were truly those of a state without any binding to C69, by examining the nitrile frequencies of the double mutant [C69A, F96oCNF]. Using the same procedure as before, we incubated the cells expressing the [C69A, F96oCNF] PYP with pCA before extracting the PYP using a Ni-NTA affinity column and compared the nitrile frequency observed by FTIR to the *apo*-protein F96oCNF and pCA-bound F96oCNF (no mutation to C69). We chose F96oCNF as the probe for this test because there is a shift upon pCA-binding that appears robust both inside and outside of the cell. The FTIR results show a clear similarity of the *apo*-protein F96oCNF to the [C69A, F96oCNF] mutant incubated with pCA (Fig. S18, Table S12), verifying that the pCA is modifying C69 inside the cells and that removal of the cysteine prevents a binding interaction from taking place.

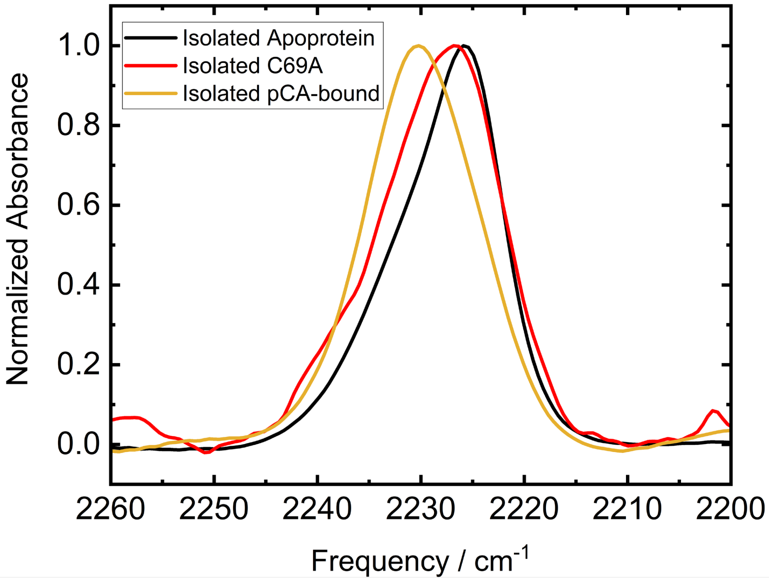


**Figure S18.** Comparison of FTIR spectra of isolated F96oCNF *apo*-protein (black) and isolated, pCA-bound F96oCNF (gold) to a [C69A, F96oCNF] double mutant that had been incubated with pCA while in cells (red), showing closer similarity between C69A and the *apo*-protein.

**Table S12.** Frequency Analysis, [C69A F96oCNF] PYP variant using FTIR.

| **Frequency Analysis** | ***Apo*-PYP** $\bar{\nu}_{Nitrile}$ **/ cm$^{-1}$** | ***Apo*-PYP Width / cm$^{-1}$** |
|---|---|---|
| **Spectral Fitting** | 2228.0 | 13.7 |
| **Peak Maximum** | 2226.7 | ---- |

### Section S5: Activation of p-Coumaric Acid

**a. Chemical reaction of pCA with carbonyldiimidazole:** Having established the capability to observe nitrile vibrational probes at low concentrations, we next sought to demonstrate a proof-of-concept study for using these nitrile reporters to detect the binding of drug-like molecules within live cells. PYP from *Halorhodospira halophila* is a well-characterized system known to form a covalent adduct with its chromophore pCA through a nucleophilic attack by the thiol sulfur of the cysteine residue at position 69 (Cys-69) within its binding pocket. In the native organism, this reaction is carried out enzymatically;[18,19] when expressing PYP recombinantly, the pCA is typically incorporated synthetically. In this process, the carboxylic acid motif of pCA is first activated to form a strong electrophile,[3,20] effectively mimicking a covalent drug bearing an electrophilic "warhead" that is highly susceptible to nucleophilic attack. This electrophilic center at the carbonyl is specifically designed to undergo a rapid nucleophilic addition reaction with the highly reactive thiol side chain of the Cys-69 residue within the *apo*-PYP binding pocket, leading to the formation of a stable covalent thioester linkage.

Our activation procedure involves reacting pCA (Sigma-Aldrich) with carbonyl diimidazole (CDI, Sigma-Aldrich) to yield an acyl imidazolide "warhead." This reaction follows the Staab reaction scheme and has been reported in several prior studies, more recently by Foreo-Doria et al.[21] in their reaction Scheme 2. Consistent with their findings (Scheme 2 in their main text, Product 3b), our procedure uses an approximately 1:1.1 molar ratio of pCA to CDI. Specifically, 1.54 mmol (253.3 mg) of solid pCA, which possesses a carboxylic acid functional group, was combined with 1.72 mmol (279.5 mg) of solid CDI in a rigorously dried 250 mL round-bottom flask. A dry stir bar was included to ensure efficient mixing throughout the reaction.

Reaction of carboxylic acids with CDI typically occurs in two steps: nucleophilic attack of the CDI carbonyl by the deprotonated pCA carboxylate to yield an acid anhydride, followed by nucleophilic attack of the anhydride by the liberated imidazole to generate an acyl imidazolide, plus free imidazole and carbon dioxide (Fig. S19).[22]

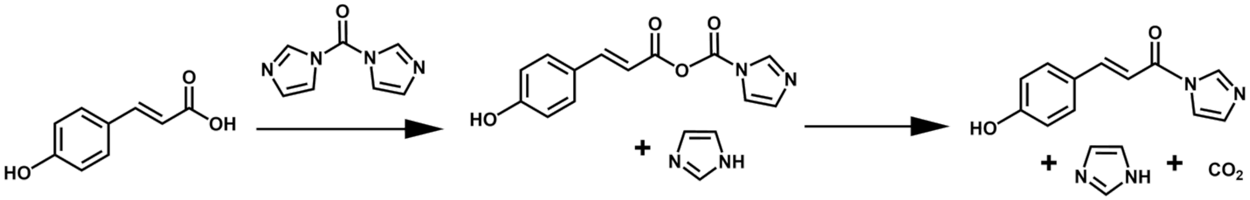

**Figure S19.** Anticipated reaction scheme for activation of p-coumaric acid by carbonyl diimidazole (CDI).

To create an inert atmosphere necessary for moisture-sensitive reagents and intermediates, the flask was sealed with a rubber septum and thoroughly purged with argon gas for 30-60 minutes. This displacement of atmospheric air, particularly water vapor and oxygen, prevents unwanted side reactions that could diminish the yield and purity of the desired product. Following the argon purge, approximately 24 mL of anhydrous tetrahydrofuran (THF, Sigma-Aldrich) was carefully added to the flask while the mixture was being stirred rapidly. Anhydrous conditions and absolutely dry THF are crucial as the activation reaction involving CDI is sensitive to water, which can hydrolyze the CDI (and or the activated intermediate).

The reaction between *p*-coumaric acid and carbonyldiimidazole was allowed to proceed for 1 hour at room temperature, after which the solvent was removed by rotary evaporation under reduced pressure with gentle heating (maintained at 40 °C). This mild condition ensured efficient removal of tetrahydrofuran (THF) without inducing decomposition of the typically moisture-, light-, and heat-sensitive *N*-acylimidazole product. Evaporation continued until all THF had been removed, yielding the activated *p*-coumaric acid derivative as a bright yellow oil. The identity of the product was confirmed by a combination of analytical techniques, including FTIR supported by DFT calculations (Fig. S20), $^{13}C$ NMR spectroscopy (Figs. S21-S23), and direct-injection liquid chromatography – mass spectrometry (LC-MS) in acetonitrile (Fig. S24). We note that this protocol is specific to the activation of pCA; most covalent drugs have intrinsic reactive warheads and do not require prior activation. Another difference we note is that while our covalent drug analogue must cross a bacterial cell wall, most covalent drugs target mammalian cells and thus can cross the cell membrane with greater ease.

**b. Vibrational Spectroscopy Characterization:** Density Functional Theory (DFT) calculations were carried out using the 6-311++G(2d,2p) basis set on Gaussian 16[23] and a polarizable continuum model (PCM) for dimethyl sulfoxide (DMSO) to simulate the infrared vibrational spectra of proposed reaction products (Fig. S20). The simulated IR spectrum revealed characteristic coupled vibrational modes centered around 1345 $cm^{-1}$, which are consistent with the formation of an anhydride or acylimidazole intermediate. These vibrational features are notably absent in the calculated spectra of the starting materials, CDI and pCA, suggesting a distinct spectral signature for the product species. Experimental IR spectra of the crude reaction mixture recorded in tetrahydrofuran display an absorption band at 1388 $cm^{-1}$. This band does not appear in the spectra of the individual reactants (CDI and pCA) or in pure imidazole dissolved in THF (expected in crude product mix), further supporting the formation of a new product. The correlation between the experimental and theoretical data reinforces the assignment of the observed 1388 $cm^{-1}$ feature to product-specific coupled vibrational modes between the pCA motif coupled through bonds to the imidazole motif, characteristic of either anhydrides or acylimidazolide products prompting us to further characterize this product using NMR.

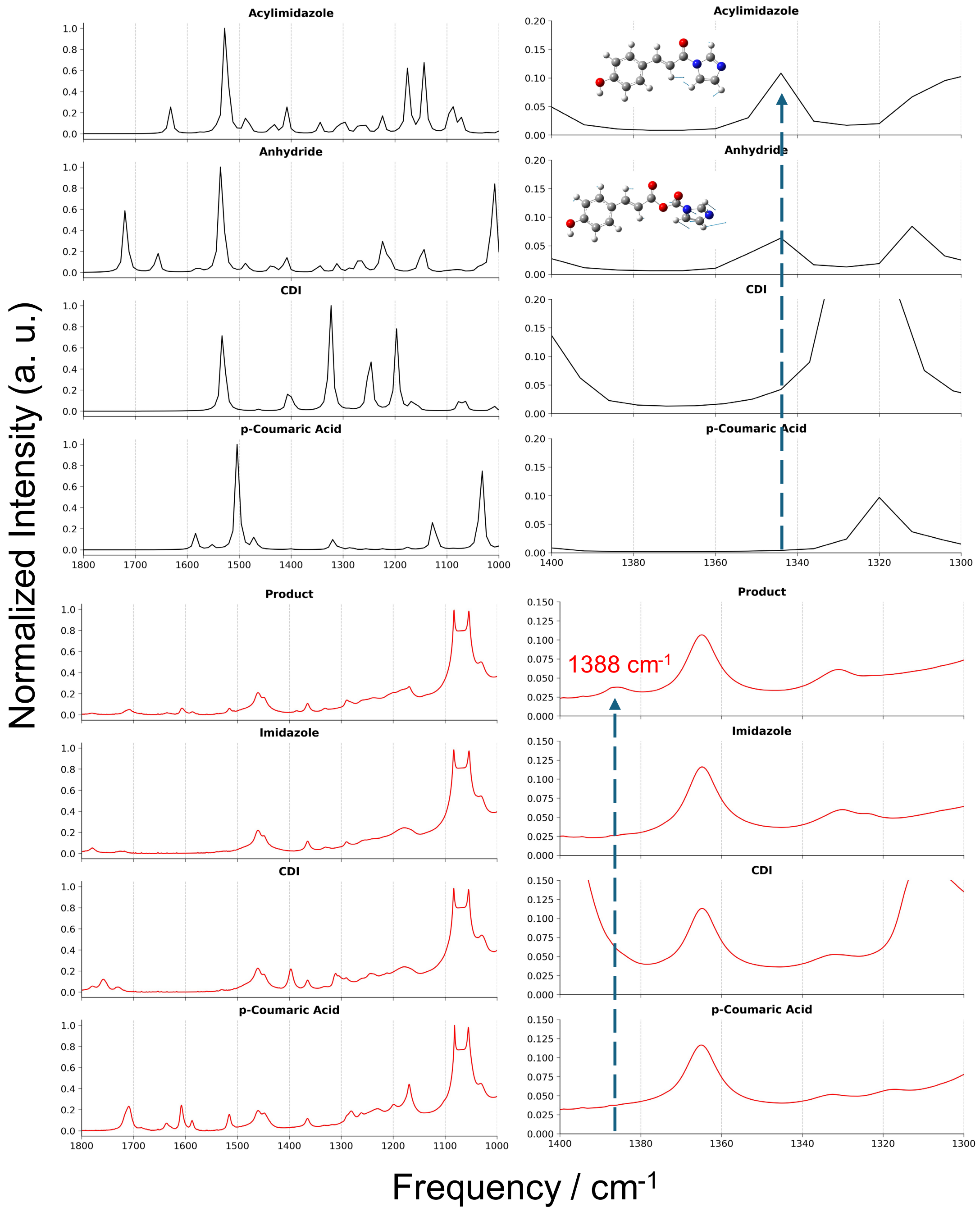


**Figure S20.** Simulated infrared (IR) spectrum (black) obtained from DFT calculations at the 6-311++G(2d,2p) level using a PCM solvent model for DMSO. The full spectrum (left) and an expanded view of the 1400–1300 cm⁻¹ fingerprint region (right) are shown. The calculations predict coupled vibrational modes near 1345 cm⁻¹, indicative of anhydride or acylimidazole formation. These modes are not observed in the computed spectra of the starting materials, CDI and pCA. Below: Experimental IR spectrum (red) of the crude product in THF, highlighting a distinct band at 1388 cm⁻¹. This feature is absent in the IR spectra of CDI, pCA, and pure imidazole, consistent with the formation of a new chemical species.

**c. $^{13}$C NMR Characterization:** We collected $^{13}$C NMR spectra of the pCA reactant and activated pCA product upon evaporation of the THF and immediate dissolving into deuterated DMSO (DMSO-$d_6$, Cambridge Isotope Laboratories, Tewksbury, MA; 256 scans 400 MHz Varian NMR). In all cases, spectra were referenced with respect to the DMSO solvent peak at 39.52 ppm. We first considered the $^{13}$C NMR spectrum of the (non-activated) pCA reactant in DMSO-$d_6$ (Fig. S21) and assigned with reference to a previously reported and assigned spectrum[24] as well as an online NMR prediction algorithm.[25] The activated pCA product spectrum (Fig. S22) broadly looks like the unreacted pCA spectrum, with some key changes. First, new peaks at 135.6 ppm and 121.8 ppm reflect the presence of imidazole and imidazolide adducts. Two of the previously observed peaks have noticeable shifts, including the carbonyl (g) which shifts 1.5 ppm downfield, and the α-carbon (f), which shifts about 1.8 ppm downfield, indicating that these nuclei are in the closest proximity to a chemical change involving the carboxylic acid. Indeed, the peaks that we observe in our spectrum are practically identical to those observed for a previously reported $^{13}$C NMR spectrum of the same activated product by Forero-Doria, et al.[21] Their purified product, though in a different solvent (methanol-$d_4$), shows the same number of $^{13}$C peaks as ours, but with smaller peaks corresponding to two imidazolide carbon types given that free imidazole had been removed using column chromatography. These imidazolide carbon peaks appear to overlap significantly with the free imidazole peaks so that only one set of peaks is observed in our spectrum. Another consistency is the extremely small peak corresponding to one of the imidazolide carbons at 132 ppm, also noted as miniscule in the Forero-Doria study. Although NMR chemical shifts of the carbonyl region clearly indicate a reaction has occurred, the ambiguity associated with the imidazolide region in NMR was resolved with subsequent methods. We also note X-ray crystallographic data of an O-methyoxy derivative of activated pCA synthesized using the same procedure[26] which clearly shows the acyl imidazolide form.

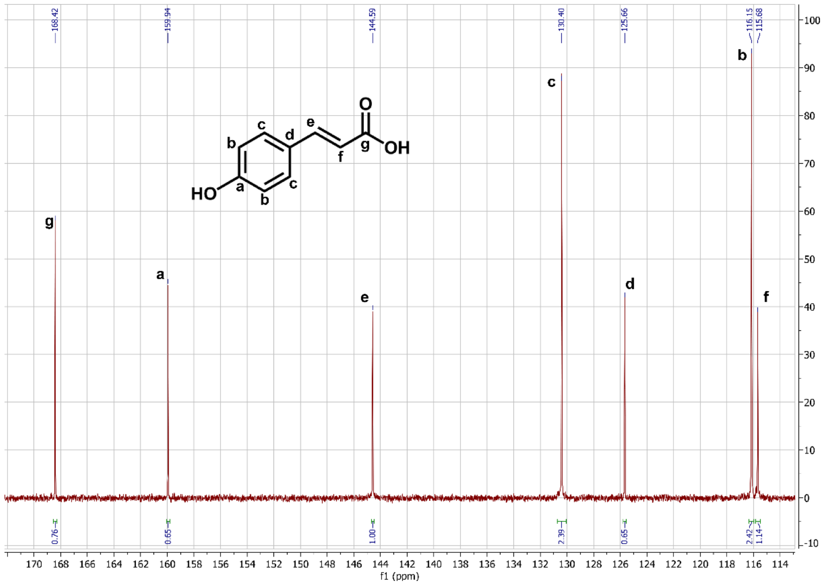


**Figure S21.** $^{13}$C NMR spectrum of unreacted p-coumaric acid in DMSO-$d_6$, with peaks assigned to carbon nuclei in the structure.

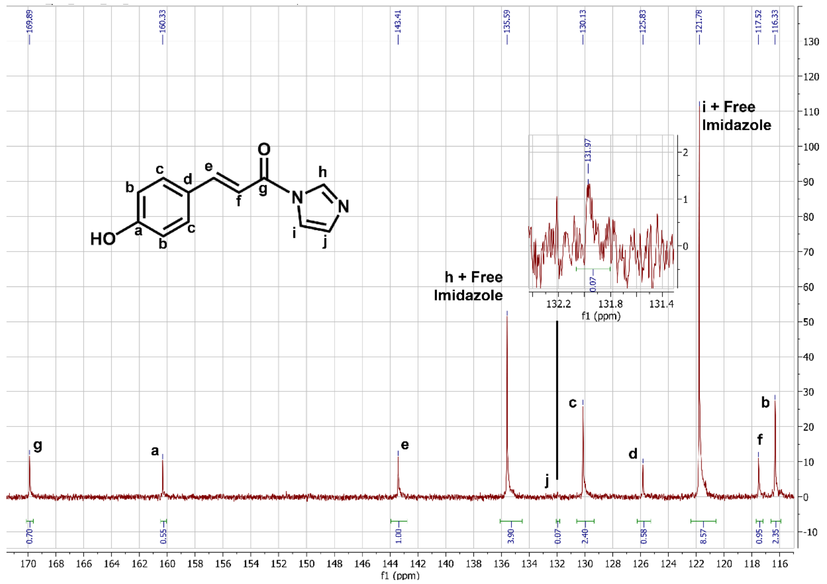


**Figure S22.** $^{13}C$ NMR spectrum of CDI-activated p-coumaric acid in DMSO-d6, with peaks assigned to carbon nuclei in the structure.

Notably, $^{13}C$ NMR analysis also indicated that the acyl imidazolide is relatively stable to hydrolysis (Fig. S23). A $^{13}C$ NMR spectrum of ~600 µl yellow oil dissolved in DMSO-*d₆* with the addition of 100 µl $H_2O$ and incubation for 1 hour at room temperature showed only minor chemical shift changes (~1 ppm or less for all carbon resonances), further confirming the stability of the acyl imidazolide under mildly aqueous conditions. These results support that the activated *p*-coumaric acid (pCA) derivative remains intact and is thus suitable for cellular uptake and specific binding to its intended biological target.

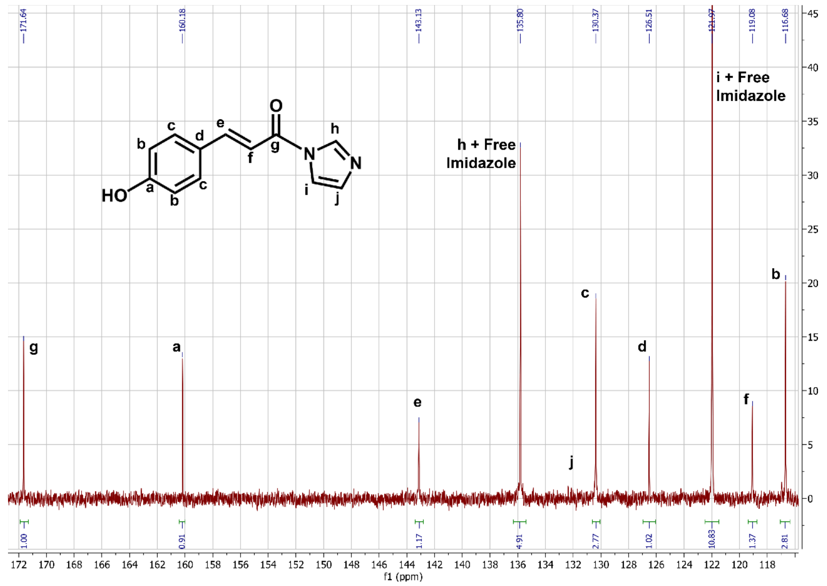


**Figure S23.** $^{13}C$ NMR spectrum of CDI-activated p-coumaric acid in DMSO-d6 after spiking in 100 µl $H_2O$ and incubation for 1 hour at room temperature, with peaks assigned to carbon nuclei in the structure.

**d. Characterization by Mass Spectrometry:** The best indication of the fully formed product is a direct-injection mass spectrum of the product in acetonitrile (Fig. S24), which clearly shows its largest peak at m/z = 215 as expected for the full acyl imidazolide product (mass=214), and a smaller peak at m/z=69 for free imidazole.

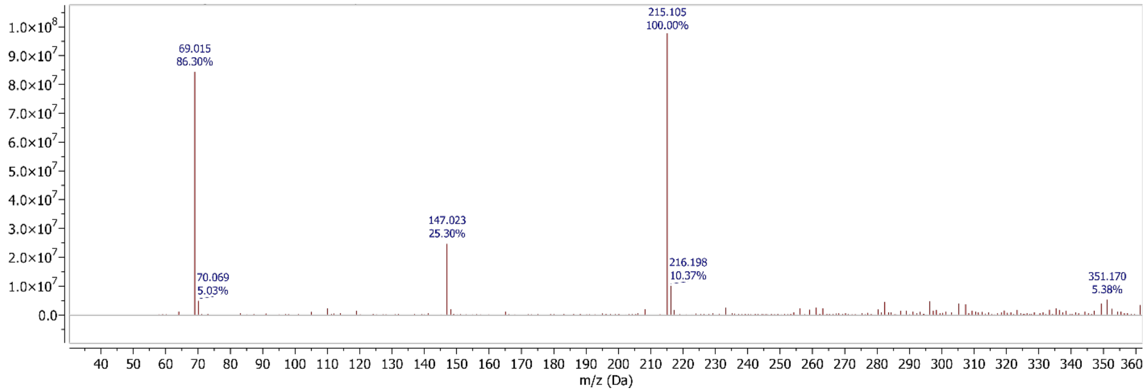


**Figure S24.** Direct-injection mass spectrum of CDI-activated p-coumaric acid in acetonitrile, using a quadrupole LC-MS with 20-mV ESI cone voltage (positive ion mode).

**e. Addition of activated pCA to PYP-expressing bacteria:** Cells were always incubated with freshly synthesized activated pCA. The molar ratio of activated pCA to *apo*-PYP variants was estimated using the expression levels as described in section S2, and adjusted to be an approximately 100-fold excess of activated pCA to each *apo*-PYP copy. Cells were added directly to the freshly evaporated yellow oil and allowed to incubate overnight at 4 °C with gentle stirring. As shown by $^{13}$C NMR results, the activated pCA is sufficiently stable in the presence of water to enable crossing of the membrane to react with PYP well before its hydrolysis. The process of cell washing and concentrating is the same with or without the incorporated pCA (Fig. S25).

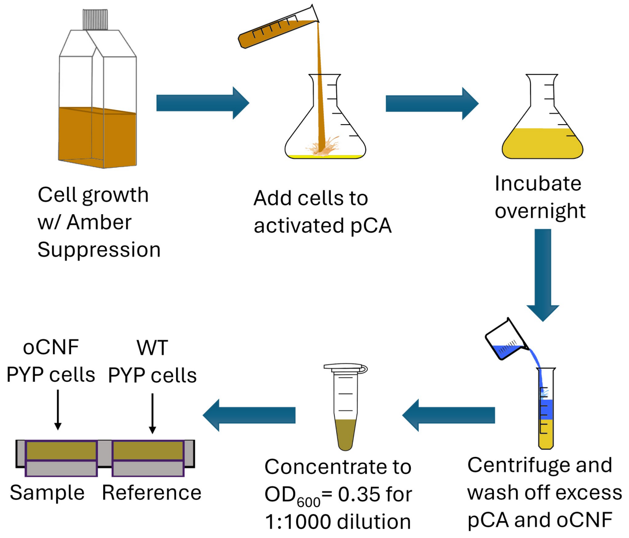


**Figure S25.** Sample preparation protocol showing the process of cell growth and optional reaction with activated pCA, prior to washing of excess oCNF and/or pCA, concentrating, and loading into a double liquid sample cell.

### Section S6: Mass Spectrometry

**a. Confirming chromophore incorporation post-lysis:** An additional test verifying the incorporation of pCA chromophore into *apo*-PYP relies on mass spectrometry, where covalent binding of pCA causes a significant and measurable mass increase. Wild-type PYP was purified from its expressing *E. coli* cells, a portion of which had been incubated overnight with activated pCA using the protocol described above. A minimal purification procedure employed here focused on Ni-column affinity chromatography to purify the His-tagged PYP, with no further tag cleaving or purification. The resulting proteins were run on LC-MS using a C8 reverse-phase chromatography column using a gradient from 0.1% formic acid in water, to methanol. Mass spectrometry was carried out using electrospray ionization with a single quadrupole detector.

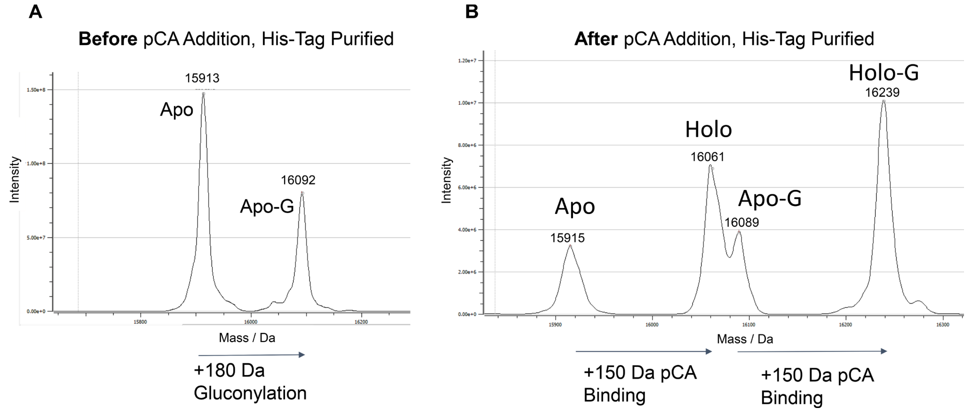


**Figure S26.** Mass spectrometry of Ni-column–purified, His-tagged PYP confirms pCA incorporation by the presence of ~150 Da mass-shifted peaks between (A) *apo*-PYP and (B) PYP from cells incubated with pCA. A fraction of the His-tagged PYP is gluconylated (*apo*-G and *holo*-G).

In both cases (with or without activated pCA incubation), the protein eluted as a single band whose mass spectrum was deconvolved to yield the major peaks shown in Fig. S26. The PYP from cells not incubated with activated pCA showed two primary mass peaks, corresponding to the His-tagged pBAD PYP (see sequence in S2) with and without a gluconylation, a common (+180 Da) post-translational modification for recombinantly expressed proteins in *E. coli* with N-terminal hexahistidine tags which we have observed previously for the tagged pBAD PYP.[27,28] The protein from the cells incubated with activated pCA showed the same peaks, plus the inclusion of larger peaks shifted by approximately 150 Da and approximately reflecting the mass increase expected due to cysteine forming a thioester with activated pCA in the chromophore binding pocket (+145 Da). Hence, mass spectrometry confirms the covalent incorporation of pCA into PYP. One aspect of note is the continued presence of *apo*-protein peaks at a lower magnitude compared to the pCA-incorporated PYP, whereas in the *in cellulo* IR measurements of the nitrile-incorporated variants, we did not see significant populations of *apo*-protein. Possible sources of this difference are (1) expression of the WT-PYP is significantly higher than the oCNF-incorporated *apo*-PYP, giving more protein in the WT cases which may not completely react with activated pCA; (2) the thioester linkage between *apo*-PYP and pCA is not completely stable in all circumstances, as detailed below, and it is possible that the acidic conditions involved in the liquid chromatography may be facilitating pCA hydrolysis on the instrument. Regardless, the ability to detect significant peaks for pCA-bound PYP confirms the covalent incorporation of chromophore detected using *in cellulo* QCL-based IR spectroscopy.

**b. Limitations of mass spectrometry methods including proteomics:** We note that while mass spectrometry is helpful here for the detection of small molecules that bind via stable covalent linkages to the target protein of interest, it may not necessarily always be useful for detection of ligands that bind weakly or noncovalently to the target protein inside cells. Even with covalent linkages such as the thioester bond between *apo*-PYP and pCA, the ligand may still be susceptible to hydrolytic cleavage under select conditions such as low pH as discussed above. We explored the use of mass spectrometry proteomics to study binding of pCA to proteins within the expressing *E. coli* cells. The most common methods involve a sample preparation that treats a cell lysate with sodium dodecyl sulfate (SDS) detergent to denature the proteins, followed by incubation with dithiothreitol (DTT) or tris(2-carboxyethyl)phosphine (TCEP) to reduce disulfide bonds and linearize the peptide sequences, capping of the cysteines using acrylamide or iodoacetamide to stabilize the linearized form, and digestion using a protease such as trypsin. We found that while the protein lost its 445-nm yellow color immediately upon denaturation by SDS at pH 7.4 (attributed to loss of the ordered chromophore pocket that stabilizes the deprotonated pCA chromophore), the chromophore was still attached by a thioester linkage at this point, giving rise to a 345-nm peak. Pure pCA at pH 8, by contrast, does not have a 345-nm peak but absorbs at 285 and 305 nm. Adding DTT or TCEP further altered the spectrum to a small 334-nm peak in addition to the larger protein 280-nm peak, and mass spectrometry showed no remaining pCA bound to PYP at this stage. Indeed, when the proteomics sample digestion procedure was applied in full to cell lysates using either DTT or TCEP, and iodoacetamide to cap the cysteines, no statistically significant hits in the *E. coli* genome were noted as being bound to pCA, not even the recombinantly expressed *apo*-PYP itself—indicating that the sample preparation scheme was harsh enough to break the thioester linkage by which activated pCA might bind itself to cysteines.

Unfortunately, the reducing conditions required for peptide linearization and mass spectrometry appear incompatible with the thioester linkage of interest in our system. While mass spectrometry-based proteomics has proven itself versatile in detecting covalent interactions including diverse electrophilic warheads attacked by cysteines, and new techniques within the field have even emerged for observing non-covalent complex formation,[29] the limitations shown by the present study indicate that challenges still remain for mass spectrometry assays involving weak covalent bonds susceptible to hydrolysis. Our method stands as an independent and entirely orthogonal, non-destructive approach to detecting interactions between target proteins and small molecules *in vivo* using the frequencies of minimally perturbative vibrational probes positioned near the binding site. Not only is our method complementary in terms of certain pitfalls characteristic to mass spectrometry methods, but it also provides new categories of information typically not available to mass spectrometry—highly specific structural details on electrostatics or hydrogen bonding patterns in the native environment, for instance.

**Section S7: Computational details**

Molecular dynamics simulations for the nitrile incorporated F28oCNF, F62oCNF, F92oCNF, and F96oCNF variants of PYP were initiated from their respective X-ray crystal structures of the *holo*-form, corresponding to PDB IDs 7SPX, 7SPW, 7SPV, and 7SJJ, respectively. The holo wild-type PYP was simulated using the crystal structure available under PDB ID 1NWZ and the chromophore under the residue HC4 was removed and the residue C69 was then protonated, while the *apo*-form of wild-type PYP was modeled using PDB ID 9O8V, a structure submitted as part of this study. Incomplete residues and sidechains were modeled using MODELLER 10.6[30] and using the Protein Analysis and Repair Server.[31]

**a. Fixed Charge Molecular Dynamics Simulations:** Fixed charge MD simulations were conducted using the AMBER99SB-ILDN force field,[32] as implemented in GROMACS 2021.3[33], on the Sherlock HPC Cluster at Stanford University. Each protein was centered in a cubic simulation box (40 Å per side), solvated with TIP3P water,[34] and neutralized with 100 mM NaCl to ensure charge neutrality and to mimic experimental ionic strength. Energy minimization was performed using the steepest descent algorithm until the maximum force dropped below 1000 kJ $mol^{-1}$ $nm^{-1}$. Equilibration was carried out under periodic boundary conditions via sequential NVT and NPT ensembles (500 ps each), with progressively relaxed restraints: initially on all heavy atoms, then only on Cα atoms, and finally with no restraints. Temperature was maintained at 300 K using a velocity-rescaling thermostat,[35] while pressure was kept at 1 bar using the Berendsen barostat[36] during equilibration and the Parrinello–Rahman barostat[37] during production. Production simulations were run under NPT conditions with a 1 fs integration time step. Non-bonded interactions were treated with a 1.2 nm cutoff, and long-range electrostatics were calculated using the Particle Mesh Ewald (PME) method.[38] All X–H bonds were constrained using the LINCS algorithm.[39]

To evaluate the effect of chromophore removal from *holo*-form crystal structures for the simulations, we compared a 100 ns simulation of the *apo*-protein (9O8V) with that of the *holo*-protein (1NWZ) following computational chromophore removal. No significant conformational differences were observed between the runs (Fig. S27). This result validated the use of *in silico* chromophore removal as a strategy for oCNF variants, for which only *holo*-form crystal structures are available. Accordingly, we applied this approach in AMOEBA simulations to evaluate local electrostatics and investigate changes linked to IR spectral shifts. We therefore proceeded by removing the chromophore from the *holo*-form crystal structures prior to running the AMOEBA simulations (Fig. S28).

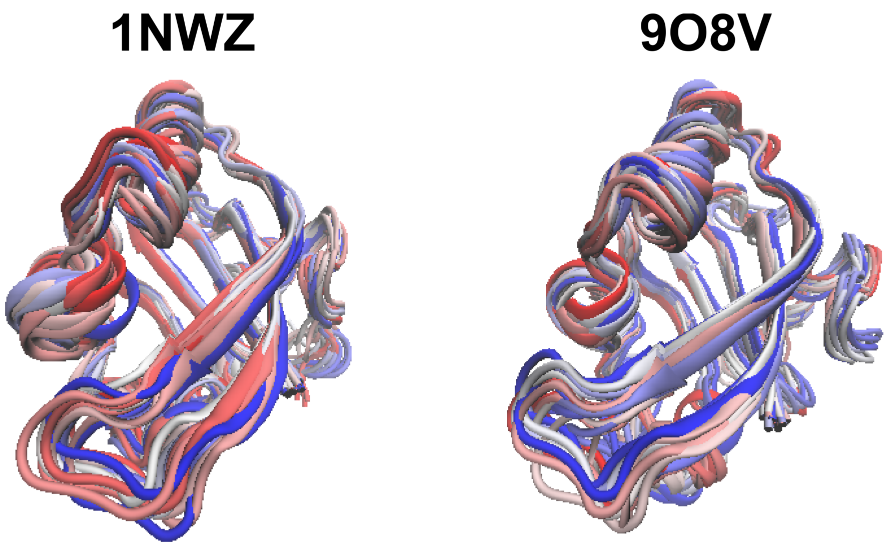


**Figure S27.** Fixed-charge molecular dynamics simulations (100 ns) were performed for *apo*-PYP starting from both the *holo* structure (PDB ID: 1NWZ, with the chromophore removed) and the *apo* structure (PDB ID: 9O8V). Snapshots were taken every 5 ns, with colors ranging from blue (early time points) to red (later time points). No major conformational changes or alterations in secondary structure were observed between the two simulations.

**b. Polarizable Force Field Molecular Dynamics Simulations:** To more accurately capture the electrostatic environment surrounding nitrile probes, we employed polarizable molecular dynamics (POL MD) simulations using the AMOEBABIO18 force field,[40] as implemented in Tinker 9.[41,42] This approach, established through extensive work by Kirsh et al.,[12] effectively incorporates polarization effects critical for modeling nitrile vibrational behavior. System setup mirrored that of the fixed-charge (FC) protocol, using a 7.5 nm cubic box containing 50 mM NaCl in water. Energy minimization was performed using the steepest descent algorithm until residual forces were below 1 $kcal \cdot mol^{-1} \cdot Å^{-1}$, with electrostatics and van der Waals cutoffs of 7 Å and 9 Å, respectively. A mutual polarization scheme with a dipole convergence threshold of 0.01 D was applied. Parameters for oCNF non-canonical amino acid incorporation have been provided earlier in Kirsh et al.[12]

Equilibration under NVT and NPT conditions followed the same restraint scheme used in the FC simulations, with the van der Waals cutoff extended to 12 Å. Simulations utilized the RESPA integrator,[43] the Bussi thermostat,[35] and a Monte Carlo barostat (300 K, 1 bar). Production simulations were run under NPT conditions with a 1 fs time step for 25 ns per replicate. Four independent runs were performed for each protein variant, yielding a total of 100 ns of aggregate trajectory per system. This was also repeated using a protonated E46 residue, but no significant changes were noted. Using Tinker, induced dipoles on the C and N atoms were recorded every 10 ps. These dipoles were divided by the corresponding atomic polarizabilities to compute the electric fields. The resulting field vectors were projected onto the –C≡N bond axis and averaged to determine the final $F_{C \equiv N}$. Data analysis scripts can be found in:

https://github.com/KozuchLab/Publications/tree/main/oCNPhe_GROMACS_TINKER

**c. Residue-Specific Fluctuation Profiles:** As previously mentioned, the resolved global structure of *apo*-PYP exhibited minimal deviation in local disorder from the *holo*-form based on Z-score analyses, maintaining structural integrity even without incorporated nitrile probes. Fixed-charge molecular dynamics simulations (100-ns duration) conducted using dual starting configurations: the chromophore-removed *holo* structure (PDB ID: 1NWZ) and native *apo* structure (PDB ID: 9O8V) also supported this observation. We also now provide residue-specific AMOEBA polarizable force field RMSF analyses for both *holo and apo* forms of the nitrile-incorporated F92oCNF and F96oCNF (Fig. S28). The *apo* system exhibited marginally higher flexibility (<1 Å RMSF increase) compared to the *holo* form, particularly near the chromophore binding pocket residues Y42, E46, and C69. This localized flexibility enhancement aligns with chemical intuition, as pCA removal eliminates stabilizing interactions in the binding cleft while preserving overall tertiary structure. The dashed traces in Figure S28 quantitatively demonstrate this differential mobility. These dynamical profiles enable us to make quantitative predictions about local electrostatic environments derived from AMOEBA-simulated hydrogen bonding patterns. The enhanced *apo*-form flexibility correlates with modified electric field magnitudes at key chromophore-interaction sites, and these computational insights directly explain the IR shifts such as the experimentally observed hypsochromic shifts upon pCA binding of F92oCNF and F96oCNF embedded nitriles, measured via QCL vibrational spectroscopy (main text Fig. 3). The combined MD-spectroscopy approach establishes a quantitative framework linking residue-specific flexibility to spectral perturbations through electric field modulation.

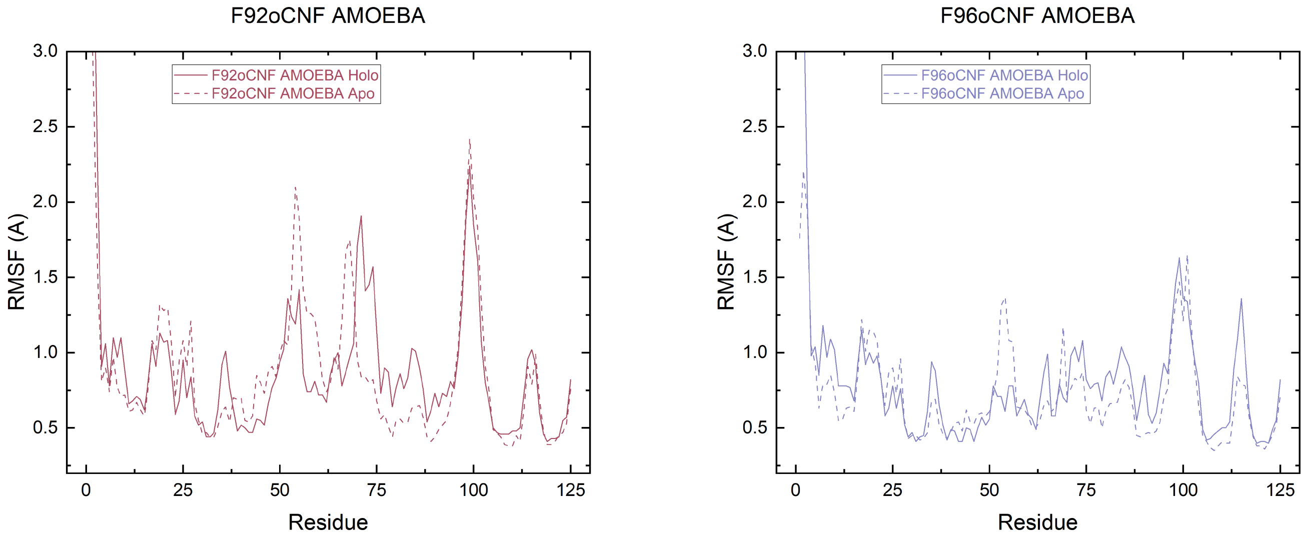


**Figure S28.** Residue-specific RMSF differences between AMOEBA simulations of F92oCNF (magenta) and F96oCNF (light blue), comparing *holo* (chromophore present, solid lines) and *apo* (computationally removed chromophore, dashed lines) forms. Only minor deviations in RMSF are observed between the *holo* and *apo* form generated by computational chromophore removal.

### Section S8: Physical interpretability of nitrile shifts

Using the simulation results, we can provide a physical interpretation for the frequency shifts of genetically encodable nitrile probes in terms of significant physical and chemical properties, i.e., hydrogen bonding and electric field. In the case of F92oCNF and F96oCNF, the variants whose nitrile frequencies shift the most upon pCA binding, we showed in the main text how these frequency shifts are dependent on hydrogen-bonding patterns involving the nitrile. These results are showcased most clearly by the AMOEBA MD simulations but are also substantiated by temperature-dependent FTIR spectroscopy involving the purified F92oCNF and F96oCNF proteins. Below, we compare FTIR spectra of the *apo*-protein F92oCNF and F96oCNF to the pCA-bound variants at different temperatures (100K and 303K), taken from the data of Kirsh et al.[12] (Fig. S29). As can be observed, hydrogen-bonded (blue-shifted) and non-hydrogen-bonded populations become more distinguishable at lower temperatures. In pCA-bound F92oCNF, the small non-hydrogen-bonding population at 100K aligns well with the *apo*-F92oCNF at room temperature, indicating that hydrogen bonding or lack thereof can help explain the frequency differences between the *apo* and *holo* (pCA-bound) forms. At warmer temperatures such as 303K, the rate of chemical exchange between the hydrogen-bonded and non-hydrogen-bonded populations becomes faster such that the two populations appear to merge in the IR spectrum. In both F92oCNF and F96oCNF, the warmer temperature leads to a peak frequency in between the hydrogen-bonded *holo* population (the primary band at low temperature) and the non-hydrogen-bonded *apo*-protein spectrum. Hence, the hydrogen-bonding explanation for the peak shifts in *apo*-PYP is in good agreement with the explanation of Kirsh et al. in the case of *holo*-PYP.[12]

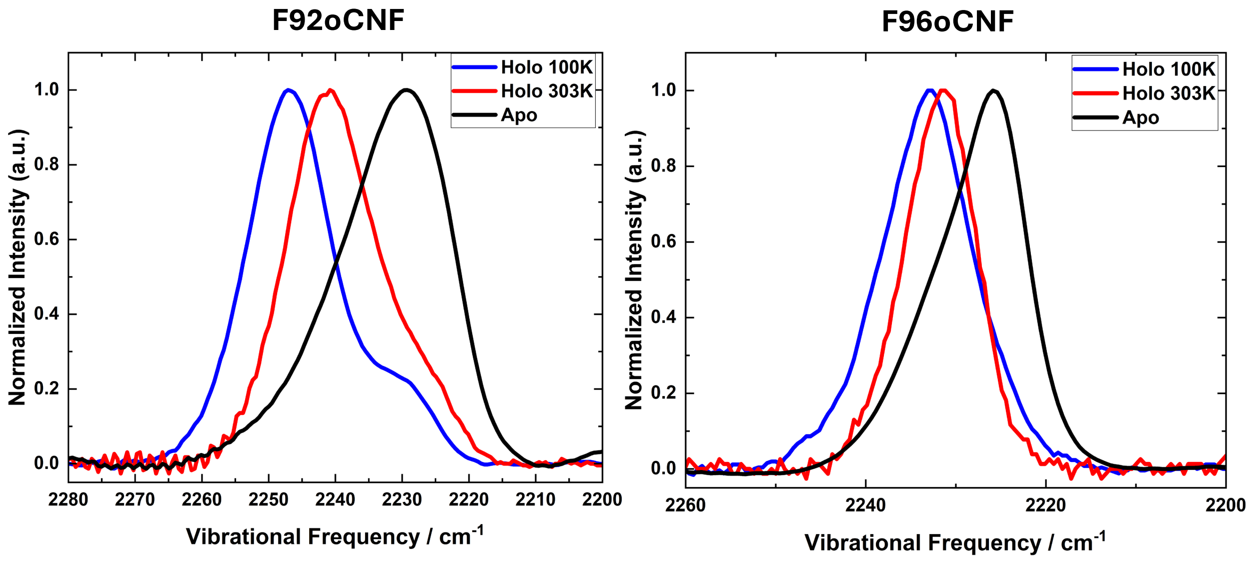


**Figure S29.** Isolated *apo*-protein F92oCNF and F96oCNF FTIR spectra are compared to FTIR spectra of the isolated *holo*-F92oCNF and *holo*-F96oCNF at various temperatures (taken from ref. 12). The low-temperature *holo*-proteins are more blue-shifted indicating a favoring of the hydrogen-bonding population at low temperatures, which his more distinguishable from the non-hydrogen-bonding population (red-shifted) that aligns more closely with the *apo*-protein.
